# Prediction of a cell-type specific mouse mesoconnectome using gene expression data

**DOI:** 10.1101/736520

**Authors:** Nestor Timonidis, Rembrandt Bakker, Paul Tiesinga

## Abstract

Reconstructing brain connectivity at sufficient resolution for computational models designed to study the biophysical mechanisms underlying cognitive processes is extremely challenging. For such a purpose, a mesoconnectome that includes laminar and cell-type specificity would be a major step forward. We analysed the ability of gene expression patterns to predict cell-type and laminar specific projection patterns and analyzed the biological context of the most predictive groups of genes. To achieve our goal, we used publicly available volumetric gene expression and connectivity data and pre-processed it for prediction by averaging across brain regions, imputing missing values and rescaling. Afterwards, we predicted the strength of axonal projections and their binary form using expression patterns of individual genes and co-expression patterns of spatial gene modules.

For predicting projection strength, we found that ridge (L2-regularized) regression had the highest cross-validated accuracy with a median *r*^2^ score of 0.54 which corresponded for binarized predictions to a median area under the ROC value of 0.89. Next, we identified 200 spatial gene modules using the dictionary learning and sparse coding approach. We found that these modules yielded predictions of comparable accuracy, with a median *r*^2^ score of 0.51. Finally, a gene ontology enrichment analysis of the most predictive gene groups resulted in significant annotations related to postsynaptic function.

Taken together, we have demonstrated a prediction pipeline that can be used to perform multimodal data integration to improve the accuracy of the predicted mesoconnectome and support other neuroscience use cases.

## 1 Introduction

A wiring diagram of the brain (connectome) is a necessary step in advancing modern neuroscience for two main reasons. First, it assists computational neuroscience by providing biologically plausible constraints on brain models and simulations (Choi and Mihalas, 2019). Second, it bridges the gap between experimental data and computational models by providing frameworks exposing its topology and other properties (Sanz-Leon et al., 2013; Ritter et al., 2013; Woodman et al., 2014). Examples of connectome based projects are the Blue Brain or the Virtual Brain that aim to create large scale cellular level models of the human brain (Markram, 2006; Markram et al., 2011; Sanz Leon et al., 2013).

The meso-scale description of the connectome (mesoconnectome) is defined at the level of anatomically distinct sub-areas within each brain region and is typically described by the use of tract-tracing invasive techniques in animal studies, or post mortem dissections in human studies (Kötter, 2007; Sporns et al., 2005; Highley et al., 1999; Lanciego and Wouterlood, 2011). The whole brain coverage provided by these techniques and the ability to delineate layer specific sub-areas make the mesoconnectome neither too coarse grained nor too spatially limited and thus suitable for developing computational models of structural brain connectivity (Oh et al., 2014; Knox et al., 2018; Betzel et al., 2015a,b).

It is difficult with tract-tracing techniques to get good whole brain coverage and they are time consuming (Sporns, 2011). As an alternative to classical neuroanatomy, genomic-based approaches have been used to describe the connectome for a number of reasons (Fornito et al., 2019). First, it is possible to infer connectivity information from genes based on the premise that postsynaptic structures have specific protein profiles and that neurons connected through synapses have highly coupled gene expression patterns (Roy et al., 2018; Sperry, (1963). Second, the recent advances in genome sequencing have resulted in gene expression data being high throughput, relatively cheap and easy to obtain (Shendure and Ji, 2008).

These advantages have led to various studies linking genomic information and structural brain connectivity with computational approaches (Baruch et al., 2008; Kaufman et al., 2006; French and Pavlidis, 2011; French et al., 2011; Wolf et al., 2011). In recent studies, a link has been established between gene expression and the mouse mesoconnectome by building predictive models and associating gene co-expression with network topology and structure (Rubinov et al., 2015; Fulcher and Fornito, 2014; Ji et al., 2014), resulting in computational frameworks for the mouse mesoconnectome.

Despite the aforementioned advances, research in the field still faces a number of limitations. Examples are the lack of cell-type specificity or synaptic density for describing specific neuronal populations in source and target brain areas that are connected through axonal projections. Such descriptions have been provided at local microcircuit level of the mouse brain but are limited to distinct brain areas such as the primary visual cortex (Lee et al., 2016). Moreover, important cytoarchitectonic features of the connectome such as the number of axonal fibers and the density of axonal arbor ending can not be extracted from models describing it as a binary network of present or absent projections between areas (Ji et al., 2014; Fulcher and Fornito, 2014).

In this work we use computational methods to measure the amount of information about axonal projection patterns present in gene expression patterns of the mouse brain and to associate them with factors related to the functional organization of genes. We have developed a pipeline, which we primarily describe in the methods section, that is available from a number of repositories. This paper is meant to describe the results we obtained with it and serve as a validation. The results section is organized as follows. First, we provide models that predict the strength or presence of axonal projection patterns given gene expression data, and we evaluate their performance on layer and cell class specific projection patterns that cover the whole brain at the mesoscale level. Second, we examine the relationship between the spatial pattern of gene co-expression modules and projection patterns in order to explain the performance of the predictive process. Third, we determine the ontological significance of predictive groups of genes in order to assess the biological relevance and causality of the predictive factors.

## 2 Methods

We built a computational framework to measure the amount of information about axonal projection patterns present in gene expression patterns of the mouse brain and to associate it with factors related to the functional organization of genes. Here we describe what data we used and how they were pre-processed as well the various steps of the analysis.

### 2.1 Materials

#### 2.1.1 Allen Mouse Brain Atlas

The gene expression data were obtained from the Allen Mouse Brain Atlas (AMBA) dataset of the Allen Institute for Brain Science (table 1), (Lein et al., 2007). The in situ hybridization (ISH) technique was used to quantify ∼ 20.000 genes over multiple spatial locations from the brains of C57BL/6J (wild-type) mice which were male and 56-day-old (P56). In this technique, a probe of complementary strand of RNA labelled with fluorescent molecules binds with the RNA of dissected brain tissue.

**Table 1:**
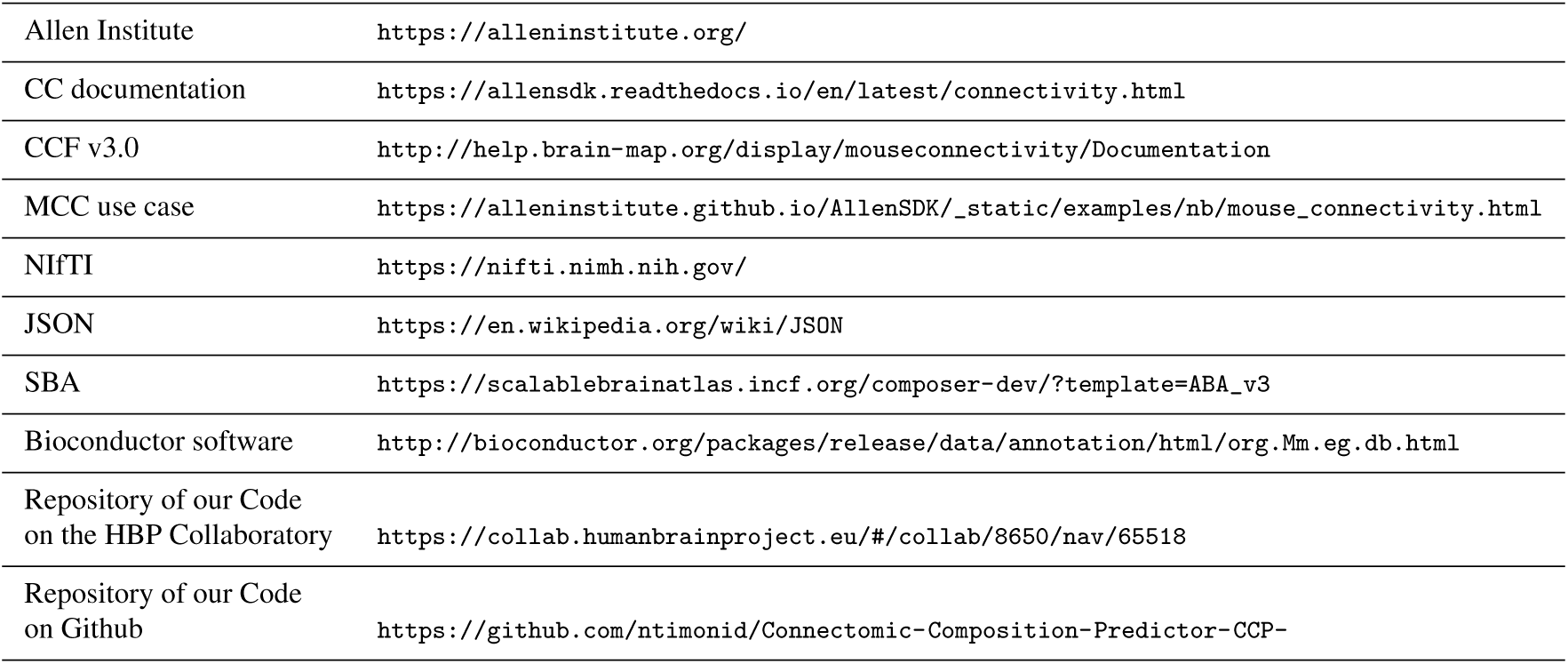
Hyperlinks for websites, tool descriptions and format descriptions related to our analysis. See main text for details.

Given that binding happens in situ, the spatial location of the gene is marked and its expression can be visualized through fluorescence microscopy (Amann and Fuchs, 2008). ISH constitutes a high throughput approach for quantifying expression energies of multiple genes in multiple spatial locations with up to 1 *µm* resolution (Lein et al., 2007).

In the study that created the AMBA dataset (Lein et al., 2007), mRNA strands were used together with fluorescence microscopy in order to visualize the gene expression energy. The result of this analysis was a set of sagittal and coronal brain slice images containing the expression energy of ∼ 20000 and ∼ 3300 individual genes respectively (Lein et al., 2007). In our analysis, the coronal slices were selected because their in plane resolution was higher.

#### 2.1.2 Allen Mouse Brain Connectivity Atlas

The axonal projection data were obtained from the Allen Mouse Brain Connectivity Atlas (AMBCA) dataset. These data were based on the anterograde tract-tracing technique that was used to quantify the strength of axonal projections within the brains of P56 wild-type and transgenic cre-line mice. In anterograde tract-tracing, fluorescent molecules are injected to a source brain area and they reach target brain areas by being transported along the axons and reaching the axonal terminals (Oh et al., 2014; Harris et al., 2018).

The AMBCA dataset is comprised of ∼ 1400 anterograde tract-tracing experiments, for which the projection density was quantified using two-photon microscopy. There were 14 major transgenic cre-lines used that resulted in the expression of label according to different laminar profiles and different cell classes within each cortical area (Harris et al., 2014, 2018). The cre-lines together with the wild-type data constituted the 15 tract-tracing categories processed in this study and in their raw form consisted of brain slice images containing projection densities of multiple target brain areas at 1 *µm* resolution (Oh et al., 2014; Harris et al., 2018).

#### 2.1.3 Allen Pre-processing pipeline

The brain slice images were processed using the informatics processing pipeline of the Allen Institute for Brain Science (table 1). Specifically, they were registered and aligned in the same reference space according to the Common Coordinate Framework CCF v3.0 (table 1).

The end product was a 3D volumetric representation of both modalities that was covering the whole mouse brain in voxel form and was provided at 100 *µm*^3^ resolution for the projection data and at 200 *µm*^3^ for the gene expression data. Each resolution referred to the size of voxels in the 3D space and corresponded to a particular total number of voxels in that space defined by x, y and z coordinates. The total number of voxels was 132 × 80 x 114 in the 100 *µm*^3^ resolution and 67 × 41 x 58 in the 200 *µm*^3^ resolution. The last step in the informatics processing pipeline was the unionization process during which the volume of both data modalities was averaged over anatomically distinct brain areas. As a result, 2D arrays were created whose rows corresponded to brain areas and columns corresponded to tracing experiments or genes respectively (Oh et al., 2014).

#### 2.1.4 Data Acquisition

In our predictive workflow we used three sources of neuroanatomical data, namely gene expression, wild-type tracing experiments and cre-line tracing experiments, that were downloaded with the mouse connectivity cache (MCC) API (table 1).

We packaged and pre-processed the data as follows (figure 1). First, a number of experiments corresponding to the expressions of genes or tracing experiments were downloaded from the Allen Institute with the use of MCC. Second, the unionized gene expression experiments were packed in a 2-dimensional array where rows correspond to anatomical brain areas and columns correspond to individual genes. Third, for each wild-type and cre-line tracing experiment, a matrix was created with rows corresponding to brain areas and columns corresponding to individual injections associated with source brain areas. Finally, all tracing-related matrices were assembled into one aggregate data structure together with tracing-related metadata such as the cell-type and laminar specificity of injections, acronyms of source areas and injection coordinates.

**Fig. 1.**
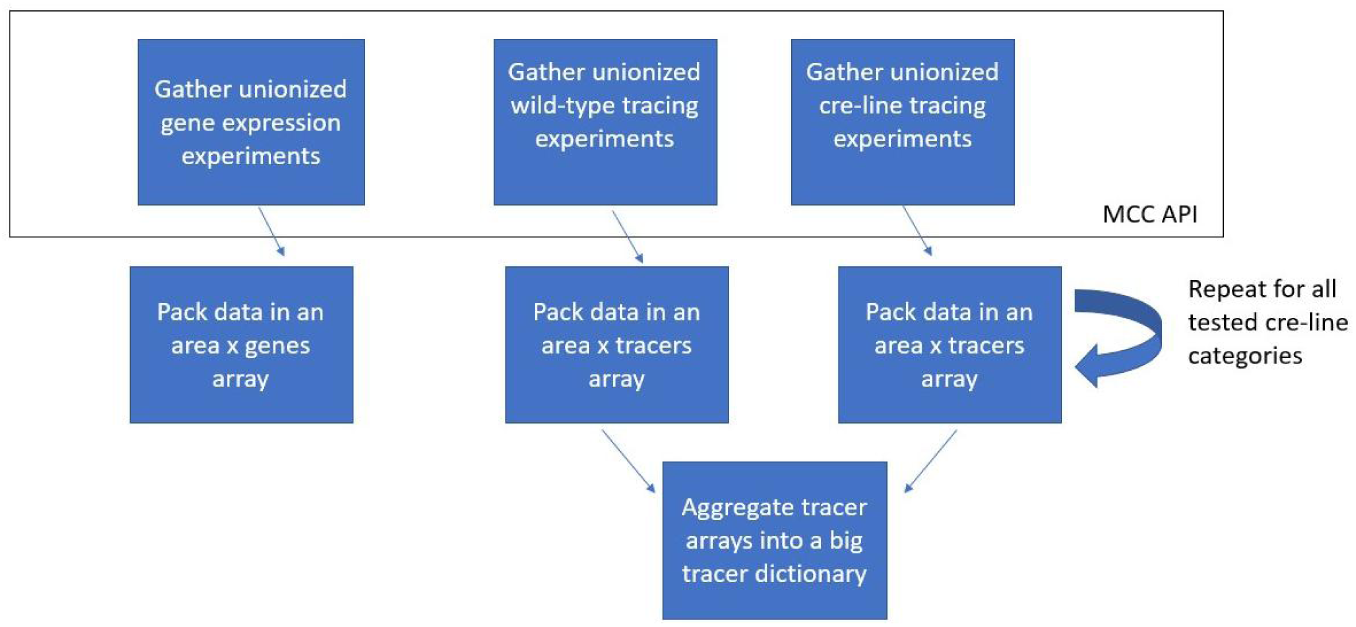

### 2.2 Procedure

#### 2.2.1 Pre-processing pipeline

We searched for not-a-number (NaN) values in the gene expression and axonal tracing datasets and removed them with a 2-step procedure based on their frequency of occurrence. Supplemental figure 1 documents that brain areas could be clustered in a group with NaN values for less than 10% of the entries, and a group with NaN values for more than 80% of the entries. We removed 610 out of 1038 anatomical brain areas defined by CCF v3.0 because they had a large fraction (*>* 80 %) of NaNs in either the gene expression or the axonal tracing datasets. The remaining NaN values in the Gene Expression dataset were imputed by taking the median value of the corresponding gene for all non-NaN brain areas (figure 2).

**Fig. 2:**
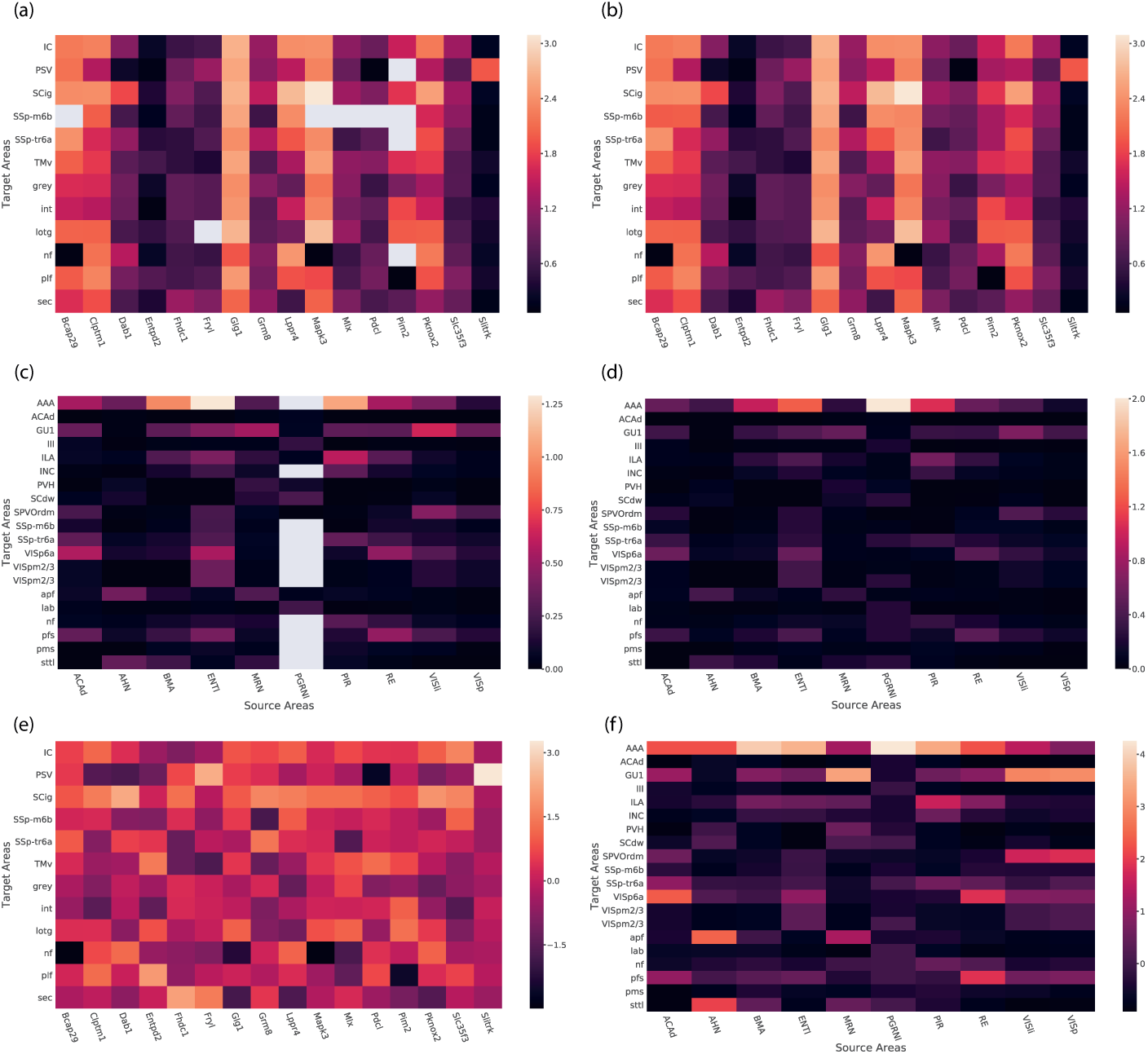
The gene expression and wild-type axonal projection datasets during different pre-processing steps. (a-b) Subset of the Gene Expression dataset containing sparse NaN values (a) and the same subset after the median imputation (b). (c-d) Subset of the wild-type projection dataset containing sparse NaN values (c) and the same subset after the sampling imputation (d). (e-f) Z-score transformation of the gene expression dataset (e) and the wild-type projection strength dataset (f). The NaN values are shown in gray. The non-NaN values have been cube-root transformed for clarity. In (a-d) the brain areas were chosen to obtain examples with some NaN values present and examples without any NaN values.

For the tracing data, a sampling-based imputation approach was followed (figure 2). To ensure that zero values would also have a chance of being used for imputing missing values, we stratified projection values per column (that is per tracing experiment) into zero and non-zero values. For each missing value present in a column, one of these groups was chosen with a probability proportional to its fraction in the non-missing data and from the chosen group a random value was drawn to be used for the imputation.

We subsequently rescaled both data modalities to obtain a proper range and distribution for use in the prediction procedure. First, a cube root transformation was applied in order to decrease the skewness of gene expression, since relative changes in expression across genes are considered to be more important than absolute ones and are usually less skewed (Ambrosius, 2007). This transformation was also applied to the axonal tracing data, since absolute changes in projection strength across tracing experiments are considered to be less important than relative ones. The cube root transformation decreased the range of gene expression values from (0 - 70) to (0 - 4) and their skewness from (ࢤ2 - 16) to (ࢤ3 - 4). This transformation also decreased the range of projection strength values from (0 - 427) to (0 - 7.5) and their skewness from (4 - 20) to (1 - 10) (supplemental figure 3). Second, z-score transformation was applied to both modalities in order to ensure that the regression-based predictive models were trained faster (Friedman et al., 2009). The z-score was obtained by subtracting the mean across areas and normalizing with the corresponding standard deviation (figure 2):

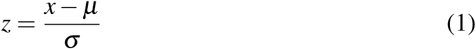

**Fig. 3:**
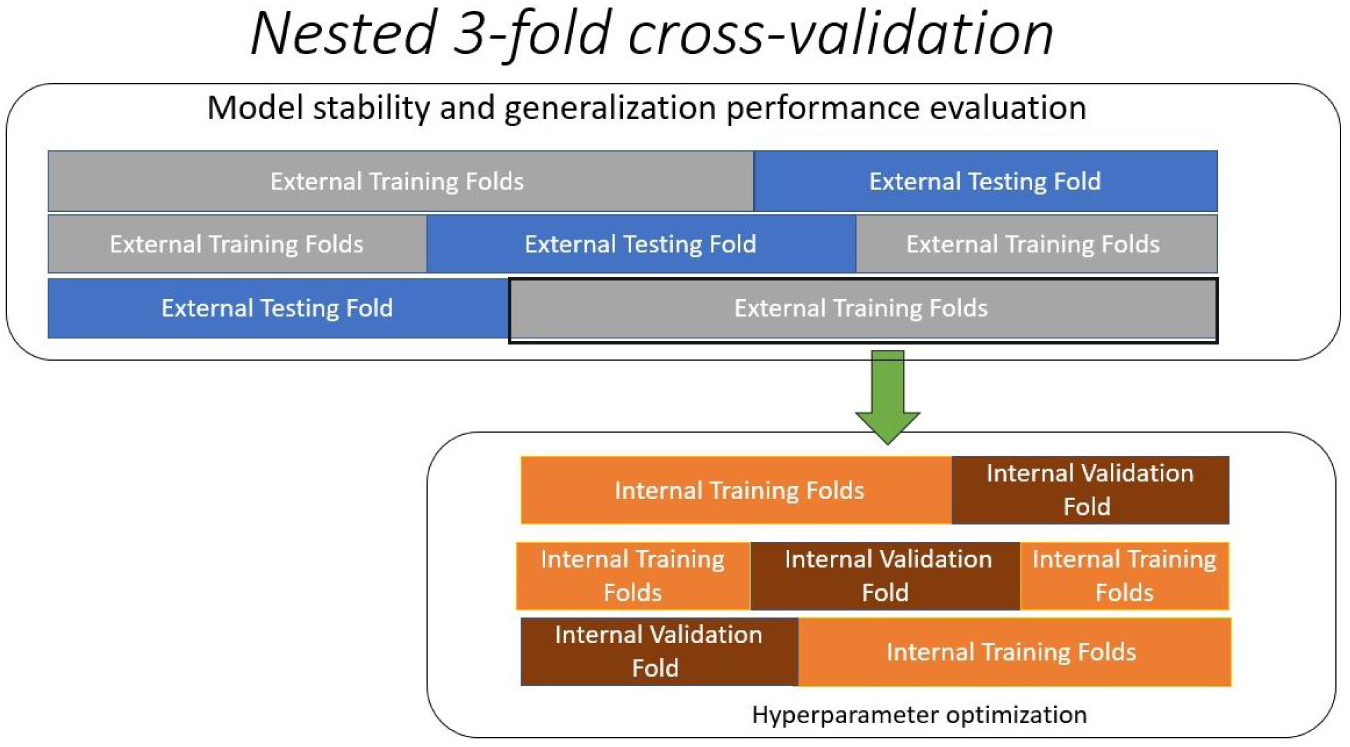
Schematic describing the structure of the nested cross-validation method.

#### 2.2.2 Model construction pipeline

A separate prediction model was built for each cre-line or wild-type category as follows. First, the gene expression data were trained with either the random forest or ridge regression method. Subsequently, model performance was validated with nested 3-fold cross-validation (Varma and Simon, 2006) and quantified by the *r*^2^ score between the measured and predicted projection patterns. The *r*^2^ score is defined as the fraction of total variance of the measured patterns that can be explained by the predicted ones (Dodge, 2008):

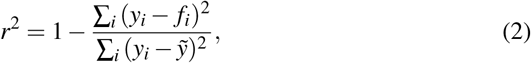

here y corresponds to a ground truth vector, 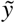 corresponds to its mean and f corresponds to the predicted version of the vector. Finally, the predicted projection patterns with their optimal hyperparameter set were extracted as model outputs.

In the paragraphs below we describe the computational methods that we used for building our models.

#### 2.2.3 Ridge Regression

Ridge Regression (also referred to as Tikhonov regularization) is a form of penalized linear regression commonly used in supervised machine learning and regression statistics (Tikhonov and Arsenin, 1977; Friedman et al., 2009). Classical linear regression fits a 2-dimensional array X to a vector y by estimating a coefficient vector w that minimizes the residuals between the actual y and the predicted *ŷ* estimated as: *ŷ*= Xw - b, where b is an intercept term.

The ordinary least squares method is used for optimizing the coefficient vector (Friedman et al., 2009):

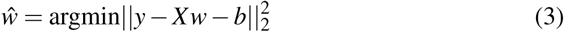

With the estimation of *ŵ*, new data *X*_*new*_ can be given as input to the model for predicting or testing *y*_*new*_:

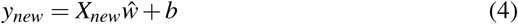

In cases of high dimensional data, where N *>* M, the dataset exhibits high variance and thus noise which hinders the generalization performance of the trained model. Ridge regression deals with the problem by constraining the size of the coefficients (Friedman et al., 2009). This is done by adding the *l*_2_ norm of the coefficients, multiplied by a shrinkage parameter *λ*, to the objective function:

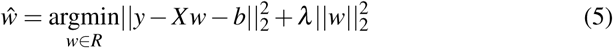

The greater the value of *λ* the greater the shrinkage of the coefficients towards zero (Friedman et al., 2009). In our analysis we utilize ridge regression in order to fit gene expression data to projection patterns of the tract-tracing data and predict unseen patterns.

#### 2.2.4 Random Forest Regressor

Random Forest Regressor is an ensemble method for performing regression tasks (Dietterich, 2000; Breiman, 2001). The basic premise for ensemble methods is that averaging reduces variance. Ensemble methods deal with high variability of predictive results by utilizing multiple models to fit and predict data, while the final result is derived by a majority voting in classification problems or by averaging in regression problems across all models participating in the ensemble (Dietterich, 2000).

In Random Forest the ensemble is comprised of multiple decision trees (Breiman, 2001). A decision tree is a directed acyclic graph or tree in which non-terminal nodes correspond to the rules of decision splits, edges correspond to each possible decision and terminal nodes correspond to the final decisions. In regression trees the internal nodes correspond to value intervals and each leaf node corresponds to the average of all training data belonging to the intervals described by its parental nodes. A decision tree is constructed by fitting training data and is evaluated by new testing data that are being assigned to a class or to values based on the decision splits of their features implied by the tree (Tan et al., 2005).

One important property of the Random Forest method is that each decision tree gets assigned a random subset of the dataset, a technique which is also referred to as bagging (Dietterich, 2000). Moreover, each decision tree can use the whole feature set of a dataset or select a random subset of it (Breiman, 2001).

In our analysis Random Forest Regressor is utilized as an ensemble alternative to ridge regression in order to investigate differences in the predictive performance between different methods. Moreover according to literature, it constitutes a robust approach against data overfitting that occurs when the training data error is significantly lower than the testing data error (Breiman, 2001).

#### 2.2.5 Dictionary Decomposition

Parallel to the predictive pipeline, the gene expression data were decomposed into transcriptional networks represented by spatial gene modules and coefficients. The Dictionary Learning and Sparse Coding method was used for decomposition, in which a data array is being represented by a linear combination of sparse but non-orthogonal modules or dictionaries and their coefficients (Mairal et al., 2010; Li et al., 2017). In dictionary learning both the coefficients and dictionaries are obtained by minimizing the deviation from the data under a L1 constraint on the coefficients (atoms) and non-negativity constraints on the elements of both the dictionaries and the coefficients:

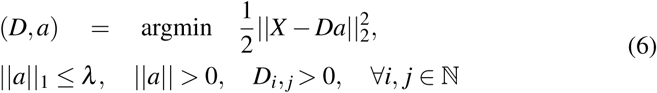

In our analysis the data array corresponded to the gene expression matrix, atoms corresponded to the coefficients of individual genes to each module and dictionaries corresponded to the spatial gene modules of the mouse brain.

There are multiple reasons to perform this gene expression decomposition. First, it allows us to visually inspect various gene co-expression patterns in the mouse brain. Second, it is a way to reduce redundancy since genes belonging to the same co-expression network have a putatively similar function across the brain (Langfelder and Horvath, 2008). Last but not least, it allows us to test multiple hypotheses regarding the effects of gene expression in predicting structural connectivity patterns. Specifically, the modification of atoms or gene coefficients can lead to altered gene expression patterns which can then be given to our models for predicting altered projection patterns.

#### 2.2.6 Internal Model Validation

For an internal evaluation of our predictive models, a technique called nested k-fold cross-validation was applied to the dataset. Before discussing the technique, it is important to first describe the classical k-fold cross-validation, from which the nested version was developed as a way to reduce bias (Varma and Simon, 2006).

A k-fold cross-validation (also referred to as k-fold CV) is a technique for measuring how well does a supervised machine learning-based model perform on new or unseen data, also referred to as generalization performance (Bishop, 2006).

In the k-fold CV method, the dataset is partitioned into k disjoint subsets of approximately equal size. D corresponds to the dataset and *D*_1_, *D*_2_,.., *D*_*k*_ are its disjoint subsets (Kohavi, 1995).

For i=1,..,k:

1. *D*_*i*_ is used as the testing set and *D \D*_*i*_ is used as the training set.
2. *D\D*_*i*_ constructs a classification/regression model using any relevant algorithm.
3. *D*_*i*_ is tested using the trained model.
4. *score*_*i*_ is estimated as the score of the model for *D*_*i*_, given any metric of interest (e.g. *r*^2^).

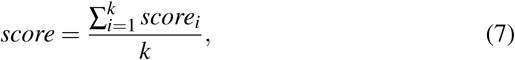

After the procedure has been completed for all k-folds, then the total cross-validation score is estimated according to eq. 7, where k is the total number of folds and *score*_*i*_ is the respective score per fold. The final score is the average over all folds (Kohavi, 1995).

Classical cross-validation is biased since both model performance evaluation and hyperparameter optimization can only be tested simultaneously on the same folds, and there is no independent set to test both factors separately. Nested cross-validation deals with the issue by nesting each training fold with internal training and testing folds and applying k-fold cross-validation to them (figure 3), (Varma and Simon, 2006). Internal testing folds are used for validating a hyperparameter set and their average predictive score (i.e. *r*^2^) is the criterion for selecting the most optimal one (figure 3, eq. 2), while external testing folds validate the generalization performance of a model. Since each external testing fold validates a model whose hyperparameter set has been selected from other folds, the aforementioned bias is avoided (figure 3, eq. 7).

Furthermore, the overall stability of the trained models can be tested by comparing the overlap of the hyperparameters selected across all external training folds. If the overlap was more than 80%, we considered the model to be stable and we trained the model on the complete dataset with the most frequently selected hyperparameter set. In this case, new data were being tested on the new complete model If the overlap was between 60% and 80% we considered the model to be moderately instable and we tested new data by averaging their predictions over all folds. If the overlap was less than 60% we considered the model to be unstable and we removed it from our set.

#### 2.2.7 Post-hoc binarization

We have provided a post-hoc approach to analyze and visualize binary projection patterns in the mouse brain, primarily for facilitating comparison to previous studies (Ji et al., 2014). Since projection strength needed to be binarized, the binarization threshold was found by maximizing the area under the “receiver operating characteristic (ROC)” curve (auROC) value (Fawcett, 2006).

For the ROC analysis, classification scores are converted to binary patterns based on a threshold and the accuracy score between measured and predicted binary patterns is estimated as the ratio between the true positive rate (TPR) and the false positive rate (FPR) (eq. 8).

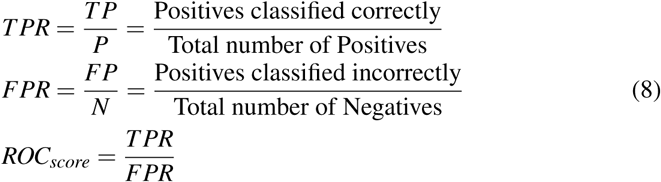

Positives and Negatives correspond to the samples from the positive and negative class respectively. It is important to mention that the terms positive and negative are conventions used to characterize binary classes in classification problems. In our case for instance, the positive and negative classes would correspond to the presence and absence of strong projections from a source area, respectively.

The strength of ROC analysis lies in the application of approximately all possible thresholds in the range 0-1, leading to a curve of approximately all possible TPR and FPR values (figure 4). The auROC is estimated as the integral of the area under the curve that represents the potential quality of classification performance under various thresholds and also reveals the optimal threshold as the point on the curve furthest away from a 45 degree line which represents the performance of a random classification (figure 4). The importance of such a line is that the actual performance quality is visible by comparing the height difference between the line and the curve: the higher the curve from the line is, the more non-random and thus significant the actual performance is considered to be (Fawcett, 2006).

**Fig. 4:**
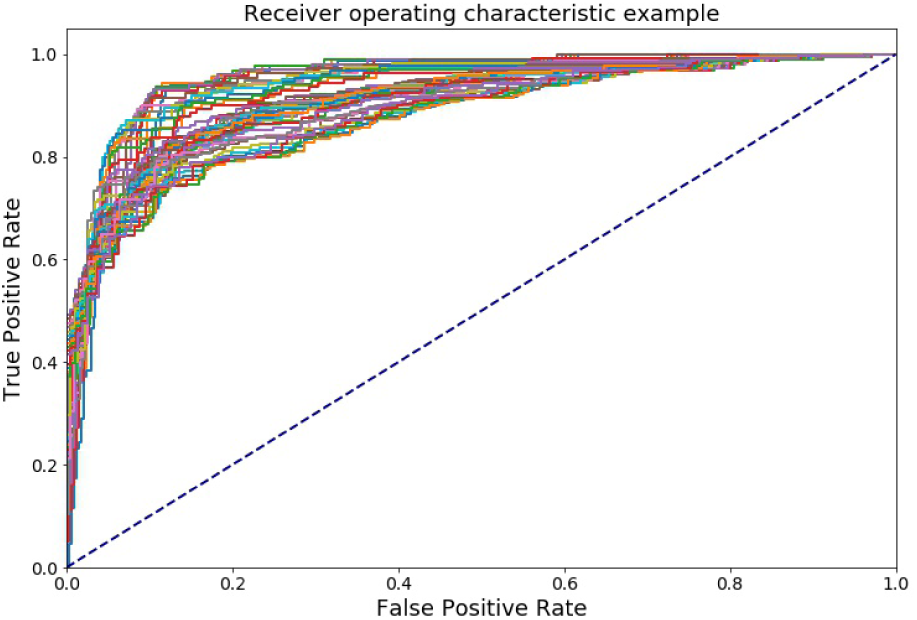
Multi-ROC curve between measured and predicted projection patterns of the Nr5a1-Cre tracing experiment. The ROC curves correspond to multiple curves induced by applying multiple thresholds to the measured data.

In order to apply ROC analysis on our continuous data, predicted patterns have to be converted to classification scores and a second threshold is needed to convert the measured projection patterns to binary ones. This is achieved by setting up an external threshold set, different from the internal one used in ROC analysis. Moreover, the predicted patterns are transformed to classification scores with the standard logistic sigmoid function 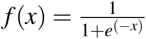

Therefore, for each external threshold in the set, the optimal auROC is estimated as the output of ROC analysis between the measured patterns binarized from the threshold and the predicted patterns that are converted to scores. The selected threshold is the one with the maximum optimal auROC value.

The Multi-ROC curve analysis in figure 4 is an example of the optimal threshold selection technique. The data corresponded to a Nr5a1-Cre tracing experiment, expressed in layer 4 and injected in the ventromedial hypothalamic nucleus (VMH). In that example, the external threshold selected was the 76th percentile of the measured data and corresponded to the curve with the maximum auROC value of 0.95. Moreover, the optimal internal threshold of the selected curve was 0.69 and was applied to the predicted data after the sigmoid transformation.

#### 2.2.8 Gene Enrichment Analysis

We used gene ontology (GO) enrichment analysis. This analysis is commonly used in the bioinformatics field for investigating the biological relevance of groups of genes (Rivals et al., 2007). In particular, the hypergeometric test that utilizes the hypergeometric distribution (Rice, 2007), was applied for estimating the statistical significance of the number of genes with a particular annotation being amongst the most predictive genes in our procedure, relative to the occurrence of genes with this annotation in similarly sized groups drawn randomly from the entire gene set (Rivals et al., 2007). We integrated GO enrichment analysis with our predictive workflow in a number of steps. First, we selected groups of genes with high coefficient scores in the prediction process (Results section, subsections 3.1 and 3.2) or genes with high coefficient scores in spatial gene modules of interest (Results section, subsection 3.4). Second, the hypergeometric test was applied to each selected gene group. Third, annotations for which the hypergeometric test returned a p-value lower than 0.05, were considered significant and were collected in a table.

The ontology annotations and the gene set for the randomly drawn subsets were taken from the *org.Mm.eg.db* database that was downloaded from the Bioconductor open source bioinformatics software (table 1) and contains genome-wide annotation for the mouse species.

#### 2.2.9 Link to Mouse Connectivity Models

We linked our predictive workflow to the Mouse Connectivity Models (MCM) tool provided by the Allen Institute for Brain Science (Knox et al., 2018). The MCM tool comprises a set of approaches, based on penalized regression, for constructing connectivity matrices on a volumetric scale of 100 *µm*^3^ or on a regionalized scale of structural brain areas. These tools enabled us to integrate the 1397 tract tracing experiments into one connectivity matrix and analyze the differences in projection patterns from different laminar profiles.

## 3 Results

We downloaded the unionized label intensities from the Allen Mouse Brain Connectivity Atlas repository (see Methods, download date = 1-07-2018) of viral tracing experiments corresponding to 1397 distinct injection sites, of which the majority (n = 498) was in wild-type subjects and the remainder were made in 14 different cre-lines of transgenic animals. Unionized means that the label intensity is averaged across all voxels belonging to a particular brain region. Here we use the common coordinate framework (CCF v3.0) to assign each voxel to a brain region. In addition, we downloaded the corresponding unionized gene expression data. The data were pre-processed to remove regions with poor quality data, impute missing values and rescale values to an appropriate range for fitting (see Methods).

### 3.1 Prediction of continuous projection strength based on gene expression patterns

We explored various fitting procedures (see Methods) for predicting the connection strength (label intensity) from the gene expression data. The two supervised learning methods used for fitting the data were random forest and ridge regression, while the performance was measured using the *r*^2^ score which represents the fraction of total variance accounted for by the model. Across all injection sites, irrespective of subject type, ridge regression based predictions yielded a median *r*^2^ of 0.54 with an interquartile range of 0.178. Random forest based predictions yielded a median *r*^2^ score of 0.42, which was lower than the one for the ridge regression based predictions (Figure 5). As an example, the data presented shown in Figure 6 were obtained using nested 3-fold cross-validation of ridge regression.

**Fig. 5:**
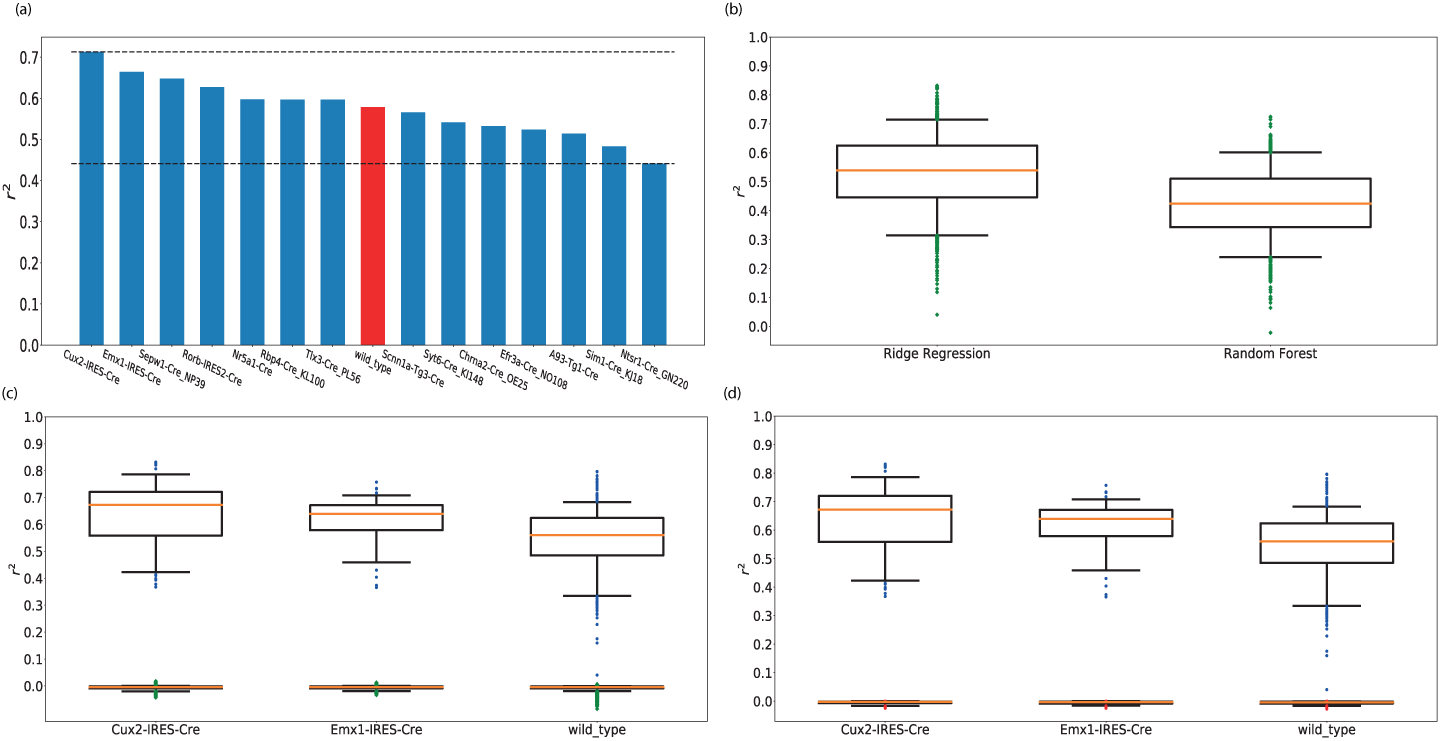
(a) Prediction scores per tract-tracing category. x-axis: tract-tracing category. y-axis: *r*^2^ values. Results for the wild-type tracing experiments are being highlighted in red in order to be differentiated from the cre-lines. (b) Comparison of ridge regression (left) with random forest (right) based models. y-axis: *r*^2^ scores. The red line is the median, the box encloses the interquartile range and the green dots are outliers which comprised 0.7 % of the injections for ridge regression and 2% for the random forest. (c) Performance comparison of surrogate (bottom panel) and actual models (top panel) for a number of tracing datasets. x-axis: datasets - Cux2-IRES-Cre (left), Exm1-IRES-Cre (middle), wild-type (right). y-axis: *r*^2^ scores. The red line and box are as in (b) while the blue and green dots are outliers for the regression of the actual and surrogate data respectively. (d) Performance comparison of null (bottom panel) and actual models (top panel) for a number of tracing datasets. x-axis: datasets - Cux2-IRES-Cre (left), Exm1-IRES-Cre (middle), wild-type (right). y-axis: *r*^2^ scores. The color conventions are as in panel (b).

**Fig. 6:**
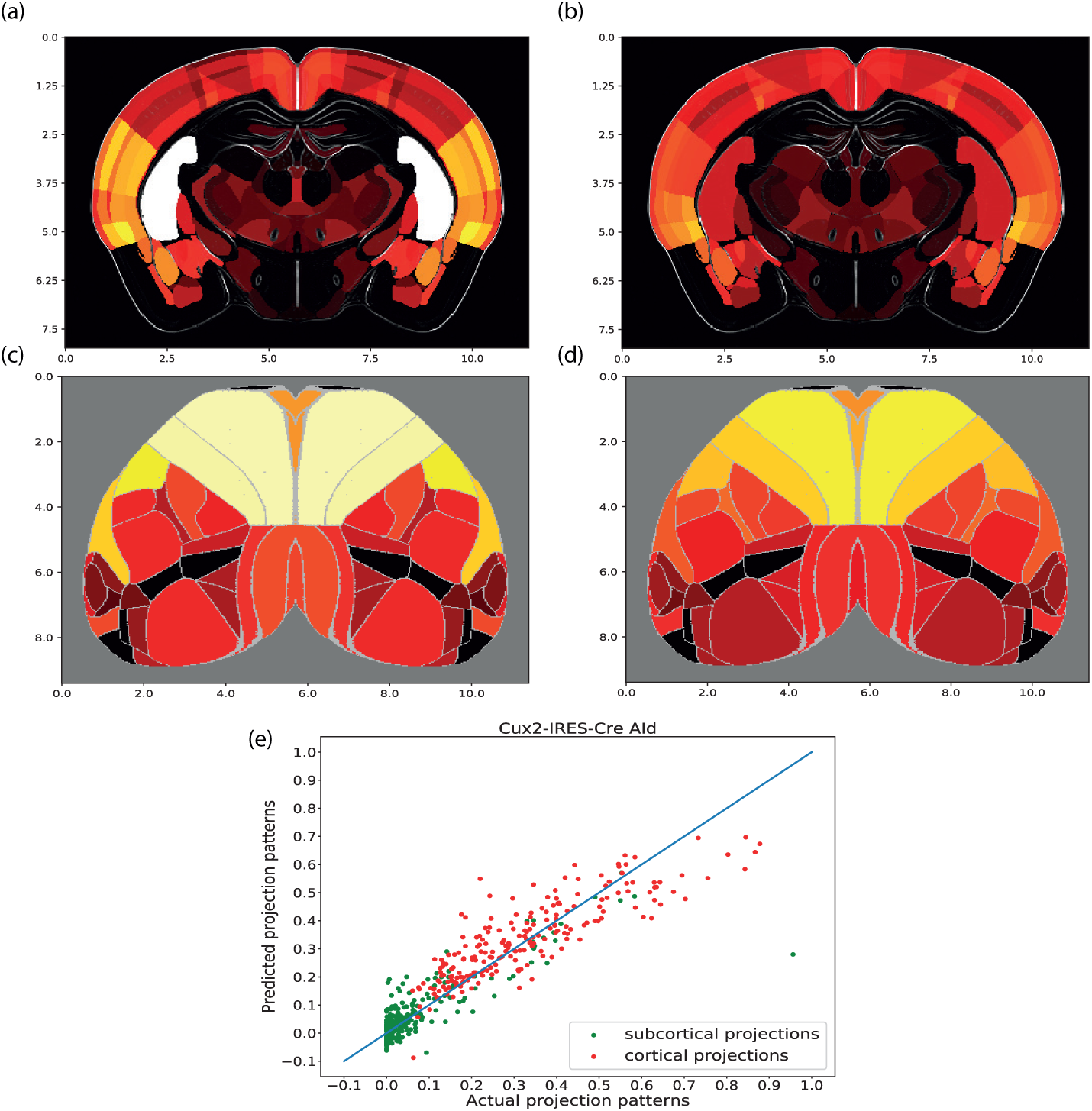
Subcortical visualizations (a,b), cortical visualizations (c,d) and a prediction performance scatter plot (e) for a Cux2-IRES-Cre tracing experiment which labeled cells in layers 2/3 and was injected in the AId area (agranular insular area, dorsal part). The *r*^2^ score for this experiment was 0.826, which was the highest score across all tracing experiments. (a,c): measured values. (b,d): predictions from gene expression patterns. The subcortical projection patterns were visualized using coronal slices of the projection volume, whereas the cortical projection patterns were projected onto a flatmap and their values have been averaged over all cortical layers. The scaling for both axes is in milimeters. The intensity of each plot was normalized by its maximum value. Cortical areas such as Retrosplenial area dorsal part (RSPd), Anteromedial visual area (VISam), trunk of primary somatosensory area (SSP tr) and Posterior auditory area (AUDpo) exhibit highly similar projection strength between their measured and predicted versions, while on the contrary subcortical similarities exist but are not as strong as the cortical ones. (e) x-axis: measured data. y-axis: predicted data. Green points correspond to subcortical projections, red points to cortical ones and the solid line is the diagonal, for which predicted values are equal to the measured ones. Cortical points are closer to the diagonal compared to subcortical ones, so they are more accurately predicted.

Variation in performance was analyzed across experiments of different tract-tracing categories. When the performance was partitioned according to transgenic cre-line and wild-type, the performance of wild-type was approximately in the center of the fit range based on transgenic animals. As the number of injections in each transgenic cre-line was much lower than available wild-type data (n = 12 to 125 for transgenic versus n = 498 for wild-type) this variation can be most likely attributed to experimental variability, rather than the specific properties of a transgenic line. Our statistical tests indicated that the difference was not statistically significant (p = 0.004 for 100 random permutations per cre-line, 14000 permutations in total, with the same distribution in set size as the cre-lines).

Predictions of projection patterns with the ridge regression-based models trained on gene expression data were significant. The ridge regression models trained with actual gene expression patterns outperformed in every case surrogate models, that were created by randomly distributing the expression intensity of each gene across areas (for three representative cases see figure 5). This process was repeated 25 times for each cre-line and wild-type tracing experiment. The predictive models that were trained with the surrogate data, also referred to as surrogate models, had a median *r*^2^ score of -0.005 and an interquartile range of 0.007 over all tracing experiments.

All of the ridge regression models outperformed the null models, that were incorporated into the analysis as an additional control (figure 5). The null models predicted unseen projection patterns by averaging values of the seen ones and thus did not account for variability across brain areas. A model was considered inaccurate when it was outperformed by those null models. The null models had a median of -0.003 and an interquartile range of 0.005 over all tracing experiments.

Predictions with low *r*^2^ values can be expected when multiple projection patterns with a noisy subset need to be predicted simultaneously. Specifically, the models were trained to fit multiple tracing experiments belonging to a particular tracing category (i.e. wild-type mice) with the same set of ∼ 3000 genes and the same hyperparameter set. In our data, 10 out of 1397 tracing experiments (0.7%) had a value in the range [0 - 0.2] for ridge regression based models, while the equivalent percentage for random forest based models was 40 out of 1397 (2.8%).

Nevertheless, performance of models with a high *r*^2^ score can be appreciated when the predicted projection patterns are visually compared with the measured ones in the form of brain slices and cortical flatmaps (for an example see figure 6).

### 3.2 Binary Predictions

Previous studies have used a binarized version of the mesoconnectome to test the accuracy of their predictive models. In order to compare our performance to these models, we developed an approach to make binary predictions as well (see Methods, figure 4). The accuracy of these predictions was quantified using an ROC analysis with as outcome the area under the ROC curve (auROC).

The median auROC value over all 1397 tracing experiments was 0.89 with a median interquartile range of 0.08 (figure 7). Moreover, performance for wild-type data matched that of the state of the art in binary projection predictions of wild-type experiments with gene expression data, such as in (Ji et al., 2014), where 93% auROC was obtained. The auROC values for all wild-type tracing experiments had a median of 0.93 and an interquartile range of 0.05. Similar values for cre-lines were obtained, which had not been subject to this analysis before (Harris et al., 2018). For instance, the auROC values for Tlx3-Cre PL56 tracing experiments, labeling cells in layers 2-6, had a median of 0.94 and an interquartile range of 0.03 (figure 7).

**Fig. 7:**
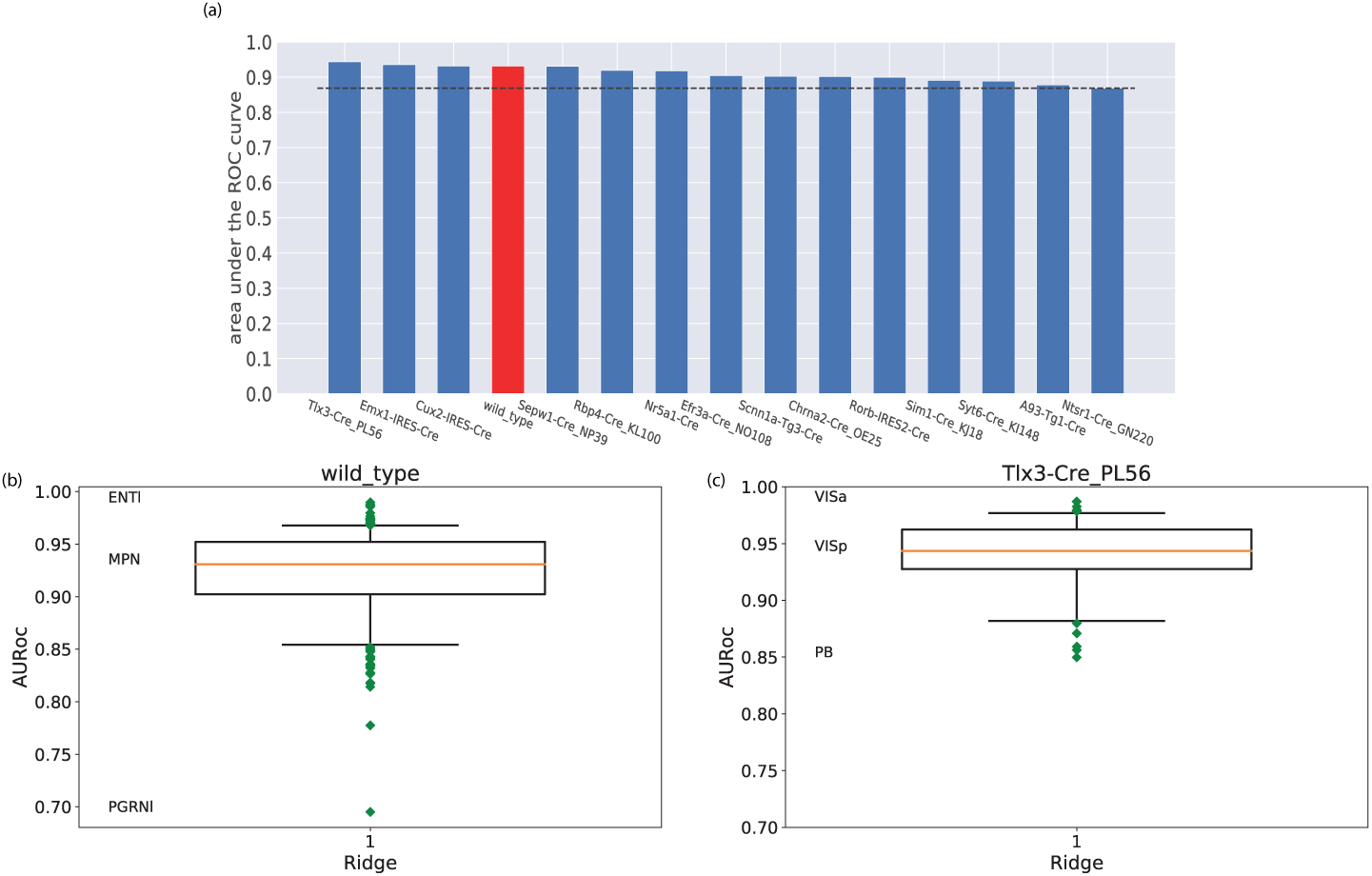
(a) Median area under the ROC curve per tract-tracing category. x-axis: tract-tracing category. y-axis: auROC values. Results for the wild-type tracing experiments are being highlighted in red in order to be differentiated from the cre-lines. (b-c) Binarized prediction performance for different categories of tracing experiments. (b) wild-type dataset. (c) Tlx3-Cre PL56 dataset. y-axis: auROC values.

Visualization of measured and predicted results, in the form of cortical flatmaps and coronal slices, allows for assessing the quality of predictions in spatial context. An example is the Cux2-IRES-Cre tracing experiment injected in AId area (figure 8), which had an auROC value of 0.98 for binary prediction.

**Fig. 8:**
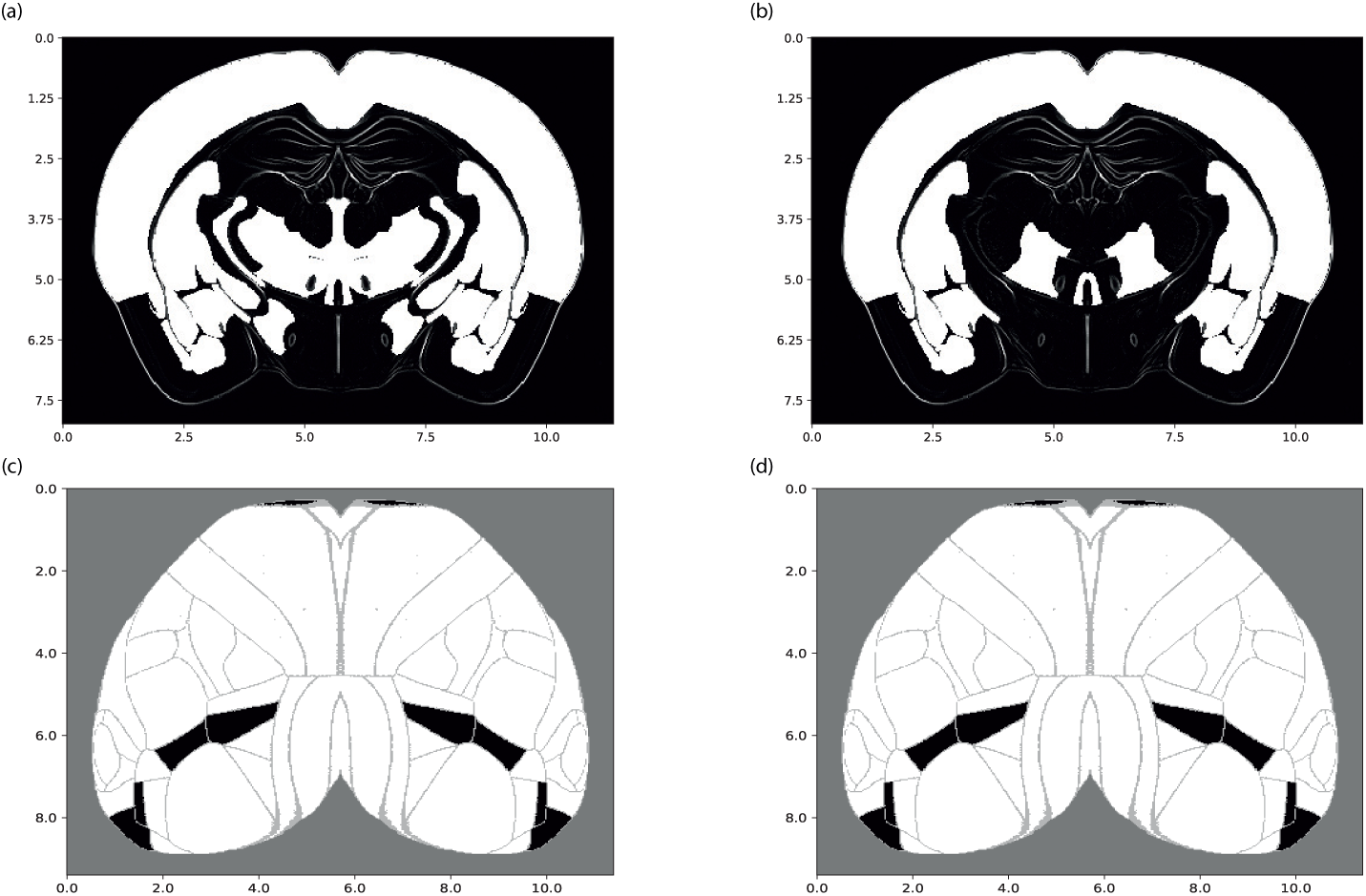
Subcortical visualizations (a,b) and cortical ones (c,d) of the binarized form for a Cux2-IRES-Cre tracing experiment injected in the AId area (agranular insular area, dorsal part). (a,c): measured values. (b,d): predictions from gene expression patterns. The subcortical projection patterns were visualized using coronal slices of the projection volume (a,b), whereas the cortical projection patterns (c,d), were projected onto a flatmap. The scaling for both axes is in milimeters. White denotes the value 1 (connections present), and black denotes the absence of a projection.

The increased performance of the models on binary predictions compared to continuous ones (figure 8) was due to reduced content of the projection patterns, which can therefore be more easily captured by the gene expression data. However, the resulting connectivity descriptions are on a very coarse-grained level which made the continuous ones more suitable for analytic purposes.

### 3.3 Creation of a regionalized connectivity array based on laminar specific projection patterns

Our predictive workflow has also incorporated regionalized connectivity models provided by the MCM tool. Specifically, we applied the MCM tool to an aggregate of the measured projection patterns from all cre-lines. The output was a laminar specific regionalized connectivity array between anatomical brain areas, for which both source and target cortical areas were laminar specific.

We investigated the differences in projection patterns across source areas with different laminar profiles. Figure 9 shows an indicative subset of the regionalized array as well as a similarity matrix between source cortical areas with different laminar profiles. The similarity matrix was created by estimating the Spearman’s rank coefficient (also referred to as rho) between the different source areas. We clustered the source areas on the similarity matrix based on their laminar profile. In addition, we applied the silhouette score for quantifying the clustering quality, which is a standard measure in clustering analysis with values ranging from -1 to 1 and reflecting the clustering cohesion (Rousseeuw, 1987). The silhouette score was 0.61, which was considered to reflect the cohesive clusters found in figure 9. Therefore, we regarded projection patterns of source areas with the same laminar profile to be more similar compared to those of different profiles.

**Fig. 9:**
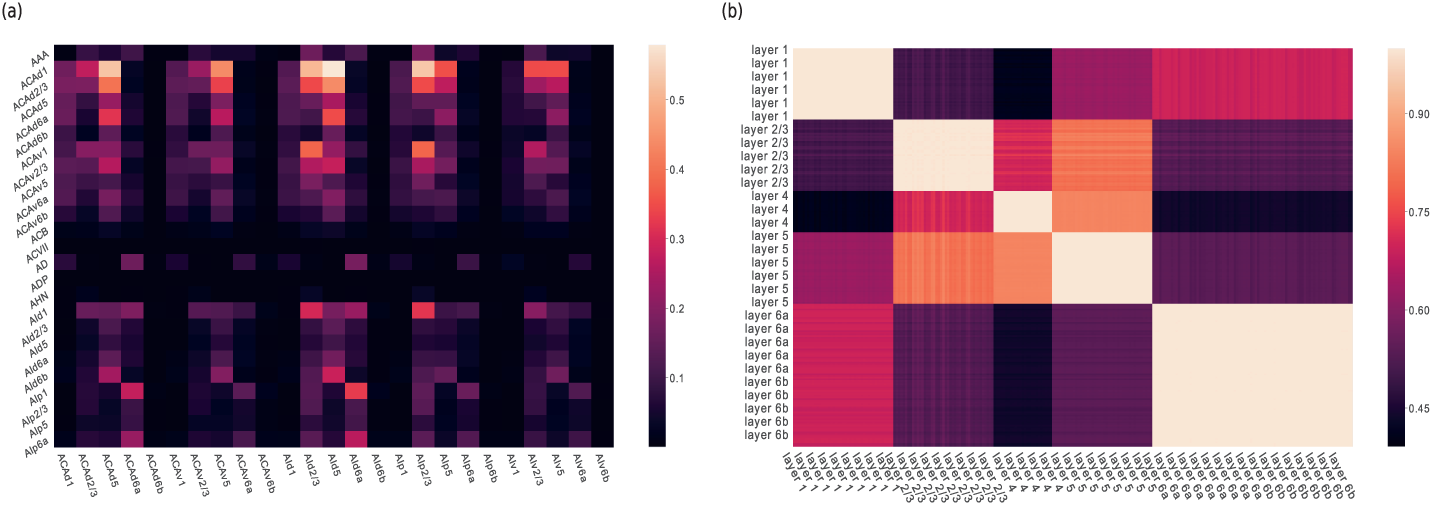
Heatmaps of a laminar specific regionalized connectivity array. (a) Subset of the array comprised from a selected set of 25 target and 25 source brain areas. x-axis: source brain areas. y-axis: target brain areas. (b) Similarity matrix of source brain areas which are clustered based on their laminar profiles. Both axes correspond to clustered laminar profiles of source brain areas. The similarity matrix was created by taking all pairs of source areas and estimating the Spearman’s rho between their projection patterns. All distinct blocks with values greater than 0.9 represent pairs of areas with the same profile. This suggests that groups of areas with the same laminar profile have more similar projection patterns compared to groups of areas with different profiles.

In order to estimate the significance of that finding, we generated surrogate clusters by randomly distributing the projection densities across source areas 1000 times. In addition, we estimated a p-value based on the number of times that the silhouette score of the surrogate clusters was greater than the score of the actual clusters. The resulting p-value was 0, which indicated that differences in projection patterns from source areas with different laminar profiles were significant.

### 3.4 Gene Module Analysis

Both the projection patterns as well as the gene expression patterns are vectors in an abstract space, for both regionalized as well as volumetric data. They can thus be written as sums of basis vectors. One can ask for a basis that most efficiently represents the variability of gene expression patterns across genes and projection patterns across injections. Therefore, we used the Dictionary Learning and Sparse Coding (DLSC) technique to find overcomplete dictionaries that account for patterns with a small number of basis vectors and small coefficients (see Methods) (Li et al., 2017). These dictionaries define groups of genes with similar spatial patterns (modules), whose spatial profile should be less noisy than the individual genes that contribute to it. Hence, in case that we do not capture the noise of individual gene expression patterns, these modules should form a better basis for predictive approaches.

With the intention of identifying functional groups with a sparse spatial distribution, we set the *λ* parameter (L1 constraint) to 1.0 (see eq. 6). The dictionary set size was chosen by training models to predict tract-tracing experiments with a different number of spatial modules, and then selecting a model with a high *r*^2^ score (see figure 10). We selected a set of 200 modules despite being second in performance (the median *r*^2^ is 0.51 for 200 modules and 0.52 for 300 modules), since the set of 300 modules was considered to be too large and their difference was considered to be an effect of variability (both interquartile ranges are 0.19 as shown by the vertical lines in figure 10, panel C). The selected dictionary set accounted on average for 10% of variability across genes and had an average spatial footprint of 88% of the brain areas. Therefore, the resulting spatial module matrix was 428 × 200, which was a significant reduction in dimensionality compared to the 428 × 3318 ISH gene expression matrix.

**Fig. 10:**
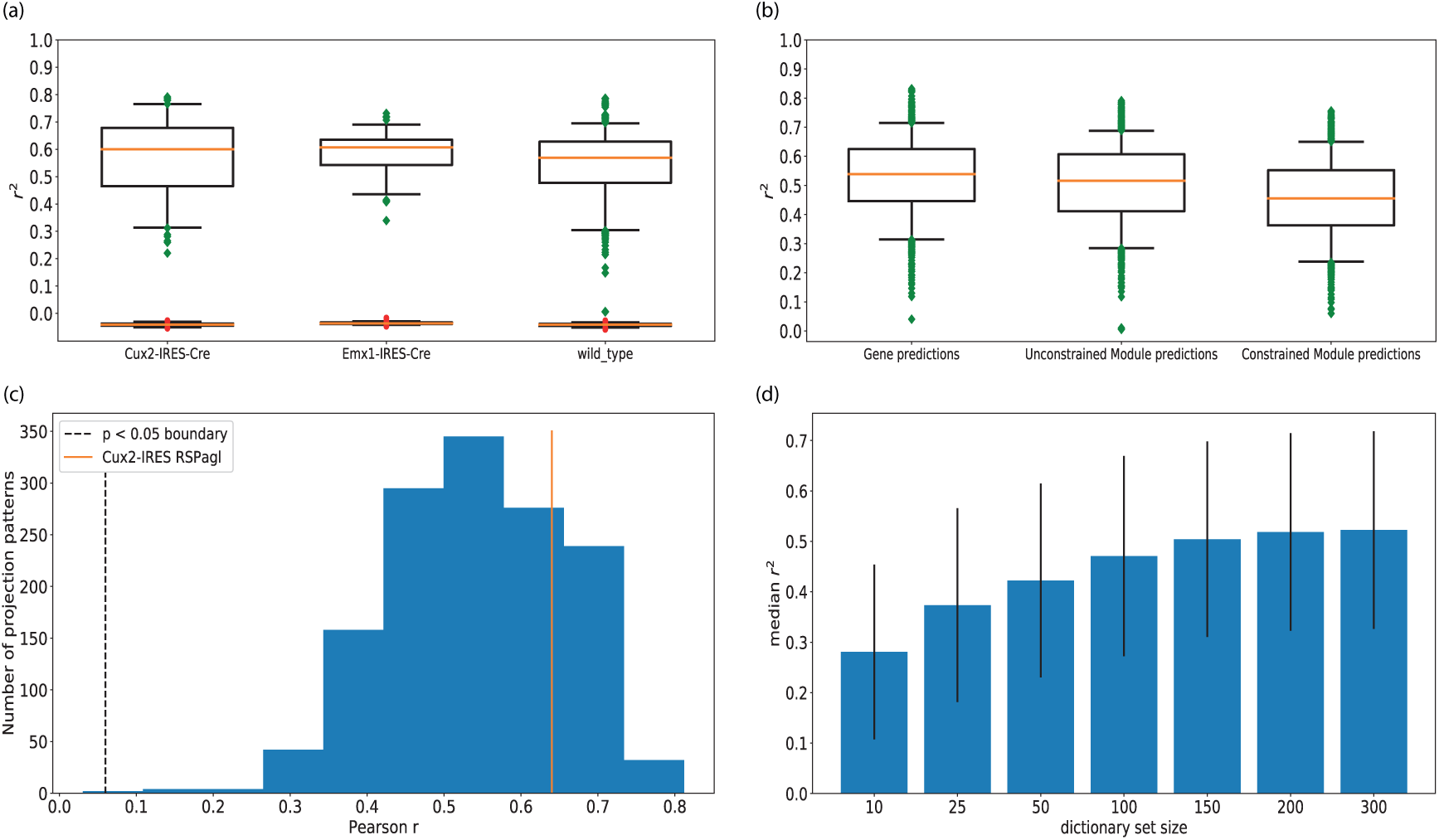
(a) Performance comparison between surrogate (bottom panel) and actual models (top panel) trained with spatial modules for a number of tracing datasets. x-axis: datasets - Cux2-IRES-Cre (left), Exm1-IRES-Cre (middle), wild-type (right). y-axis: *r*^2^ scores. (b) Comparison of predictive accuracy between models trained using spatial modules and models trained using full gene expression data. x-axis: gene expression based models (left), spatial module based models (middle) and module based models constrained with single-cell RNA sequencing data (right). y-axis: *r*^2^ scores. (c) Histogram of pearson correlation coefficients (r) between all 1397 tracing experiments and their predicted versions. The prediction of each experiment was achieved with its 3 best correlated modules as determined by pearson r. The first vertical line to the left represents the point at which all correlations left from it are not longer statistically significant (p *>* 0.05). The second vertical line to the left corresponds to the pearson r of 0.64 between a Cux2-IRES-Cre experiment injected in the RSPagl area and modules 9, 70 and 88, which is greater than the mean r of 0.54 (see figure 12). A dense distribution of correlations in the range 0.4 - 0.7 indicates that multiple spatial modules correlate with axonal projection patterns. (d) Predictive performance of module-based models with different dictionary set sizes. x-axis: dictionary set size. y-axis: median *r*^2^ score over all tract-tracing experiments. The vertical lines represent the interquartile range across the dictionary sets. The highest peak is for 300 modules with an *r*^2^ score of 0.52.

In order to examine the predictive capabilities of the spatial modules, the prediction process was repeated with models trained on the modules instead of genes. We considered an example tracing experiment for which the module based predictive model had the highest *r*^2^ score of 0.79. The tracing experiment was generated by a Cux2-IRES-Cre injection in Retrosplenial area, lateral agranular part (RSPagl). We looked for modules with the highest similarity with the projection pattern, as quantified using the pearson correlation coefficient (r). We selected three modules, labeled as 9, 70 and 88, with a pearson r of 0.51, 0.52 and 0.45 respectively. Each of these modules were non-zero in a mostly non-overlapping group of brain areas, which together cover a part of the experimental projection pattern (see figures 11,12). We analyzed the contribution of their spatial footprint in each area separately, by replacing each nonzero value by 1.0 if present in all three modules and the projection pattern, 0.8 if present in two modules and the pattern, 0.6 if present in one module and the pattern, 0.4 if present in the pattern and absent in all modules and 0.2 if present in the modules but absent in the pattern. As indicated in figure 12, there was a large overlap between the experiment and the modules in cortical areas. Subcortical areas did not have such strong coverage as cortical ones, which might be the reason why predictive performance was not higher in terms of the *r*^2^ score.

**Fig. 11:**
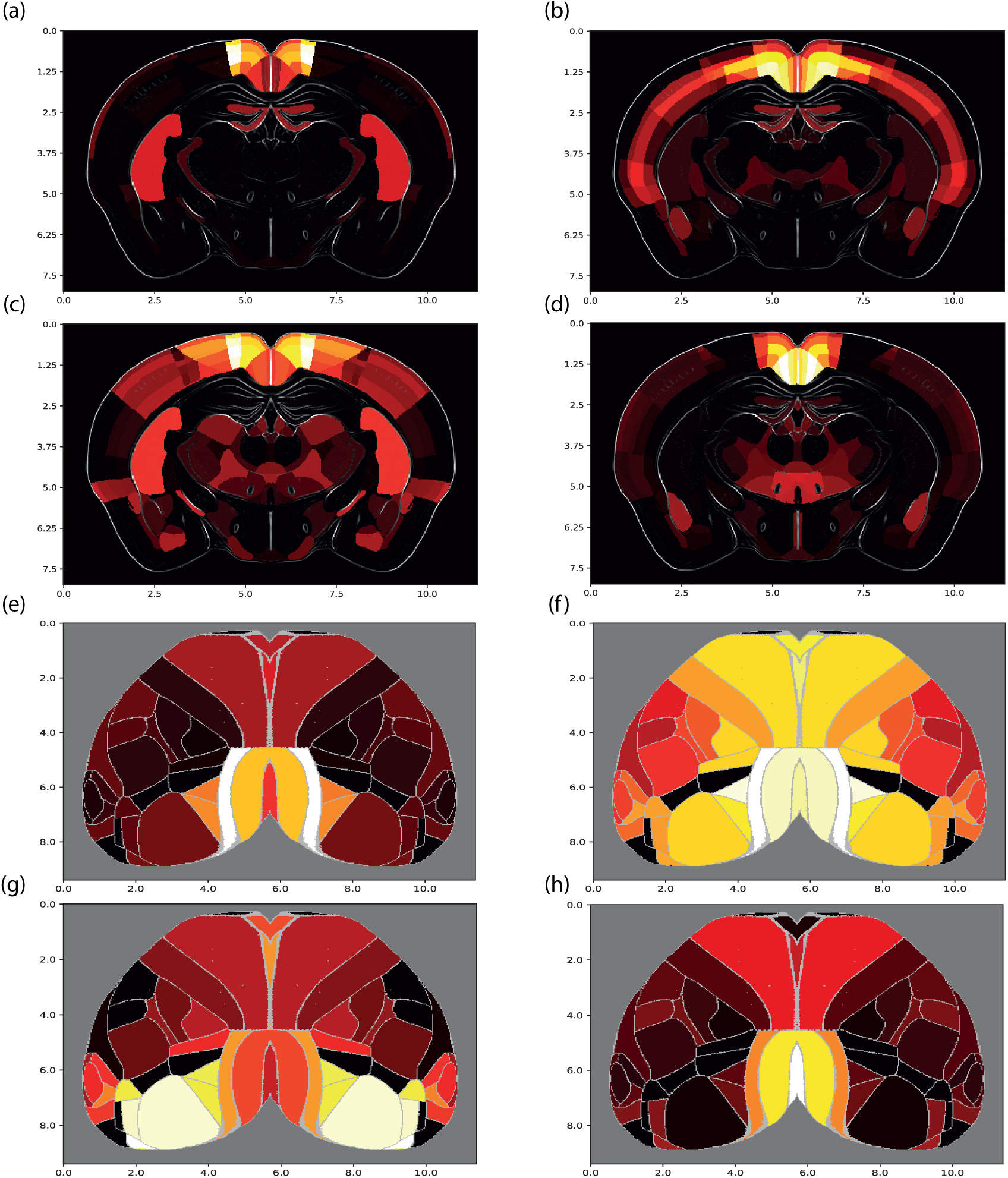
(a-d) Subcortical and cortical visualizations (a,e) for the Cux2-IRES-Cre RSPagl tracing experiment compared to subcortical and cortical visualizations of spatial gene modules 9 (b,f), 70 (c,g) and 88 (d,h). The subcortical projection patterns were visualized using coronal slices of the projection or module volume (a,b), whereas the cortical projection or module patterns (c,d), were projected onto a flatmap and their values have been averaged over all cortical layers. The intensity of each plot was normalized by its maximum value. The scaling for both axes is in milimeters.

**Fig. 12:**
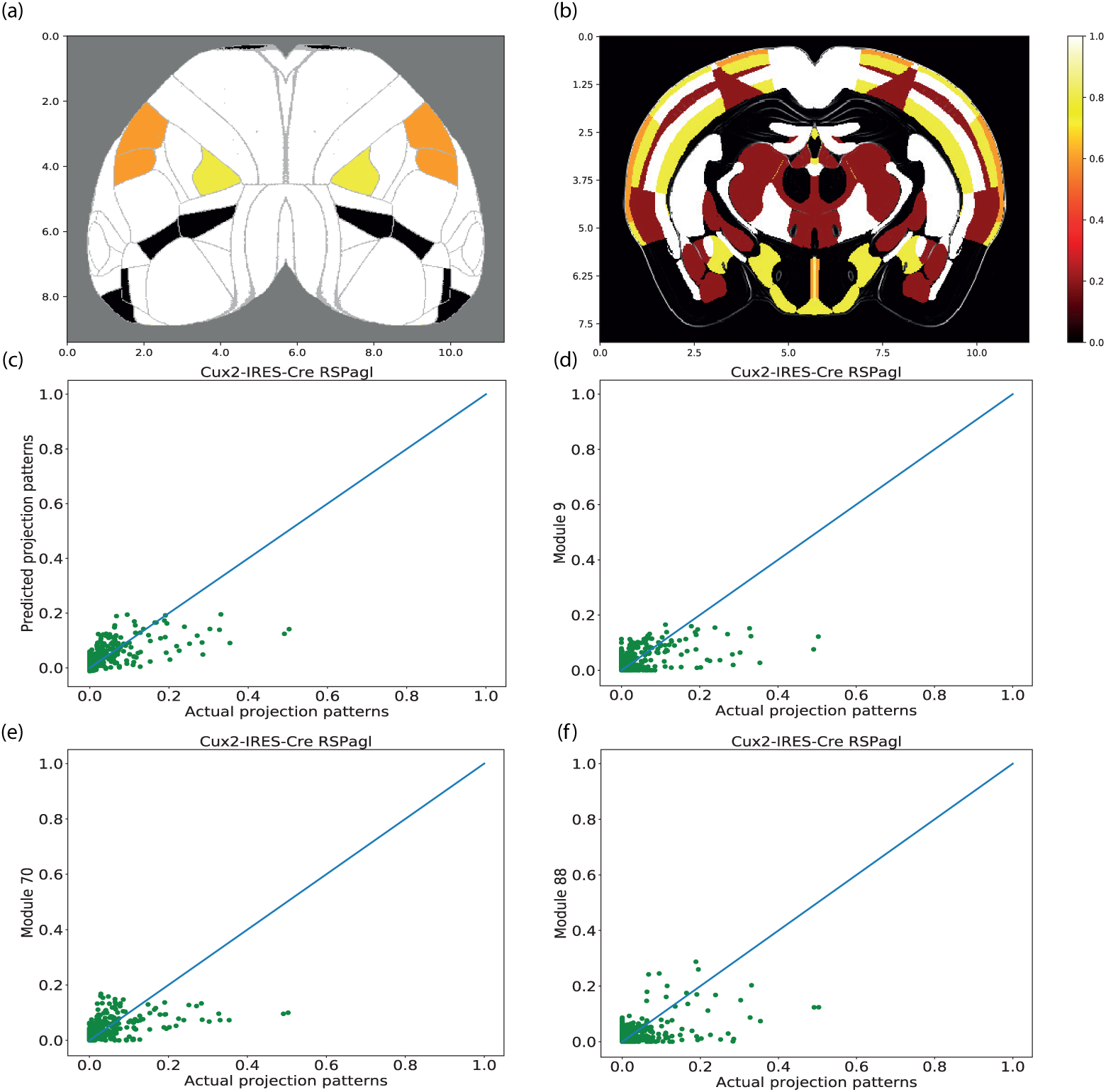
Cortical (a) and subcortical visualization (b) of a spatial footprint related to a Cux2-IRES-Cre tracing experiment injected in the retrosplenial area, lateral agranular part (RSPagl). The spatial footprint represents the overlap that exists between the RPSagl experiment and modules 9, 70 and 88 with a pearson r of 0.51, 0.52 and 0.45 respectively. Each non-zero value across brain areas is replaced by 1.0 if it was present in all three modules and the projection pattern, 0.8 if present in two modules and the pattern, 0.6 if present in one module and the pattern, 0.4 if present in the pattern and absent in all modules and 0.2 if present in the modules but absent in the pattern. The subcortical projection pattern was visualized using coronal slices of the projection volume, whereas the cortical projection was projected onto a flatmap and its values were averaged over all cortical layers. The scaling for both axes is in milimeters. There is a strong presence of white (1.0), yellow (0.8) and orange (0.6) colors, that suggests a strong overlap between the experiment and the modules and which is also reflected by a *r*^2^ score of 0.4 when the three modules are used for predicting the experiment. (c) Scatter plot between the projection pattern of the same experiment and its prediction by the 3 modules. (d-f) Similar scatter plots between the projection pattern and each module separately (d for module 9, e for module 70 and f for module 88). The solid line in each scatter plot is the diagonal, for which values across axes are equal. The scatter plots suggest that a combination of the three modules scaled by coefficients can lead to a more accurate prediction of the experiment than by the individual modules alone.

Subsequently, we calculated the pearson r between the RSPagl experiment and its prediction by the three modules. We found that this prediction yielded an *r*^2^ score of 0.4 and a pearson r value of 0.64, which was higher than the median pearson r of 0.54 over all tracing experiments. Therefore, these modules were important components to the total prediction, whereas they provided a less accurate prediction as stand-alone predictors (see figure 11,12).

This finding suggests that multiple spatial modules might be needed to reproduce projection density patterns from the mouse cortex (figures 12,10). For the predictive models trained and tested with spatial modules over all tracing experiments, the median *r*^2^ score was 0.51, the interquartile range was 0.19, and the maximum *r*^2^ score was 0.79. Therefore results were slightly lower on average than the corresponding ones for the gene predictions (figure 10). For testing the significance of module based predictions, surrogate models were built as explained in subsection 3.1 and trained with spatial modules instead of genes. All models trained for the 1397 tracing experiments had higher *r*^2^ values than the respective surrogate ones, as indicated by a number of examples in figure 10. Taken together, our findings suggest that spatial gene modules contain a large fraction of the predictive capacity of the much higher dimensional gene set.

Previous studies focused on integrating single-cell RNA sequencing with ISH data in order to provide cell-type densities (Mairal et al., 2010). Furthermore, the neuroexpresso tool has provided access to a large collection of single-cell gene expression data derived from multiple studies (Tasic et al., 2016; Mancarci et al., 2017).

In (Mancarci et al., 2017), the authors collected expression data of ∼ 11.000 genes from pooled cell-type microarray and single-cell RNA-sequencing studies. For that reason, we tested the capability of the DLSC model to provide meaningful modules by constraining it with the neuroexpresso data. We selected 74 cell-types from their repository (https://github.com/PavlidisLab/markerGeneProfile) and 2154 genes that were common between the single-cell and the ISH data, which resulted in a 2154 × 74 array of cell-type specific gene expression. Finally, the cell-type specific and the ISH gene expression arrays were given as inputs to the DLSC model that created a new array of 428 areas x 74 constrained spatial modules.

We trained the prediction models using the constrained spatial modules and evaluated the quality of the resulting projection patterns. The term unconstrained is used to describe the spatial modules from the DLSC model which was not constrained with single-cell RNA sequencing data. The median *r*^2^ score of the constrained models for all tracing experiments was 0.45, which was substantially lower compared to the unconstrained ones (median *r*^2^ of 0.51), and the interquartile range was 0.19. In addition, we created a similarity matrix between the constrained and unconstrained modules and we applied biclustering analysis to it (figure 13). The similarity matrix was created by Spearman’s rho, and the biclustering algorithm used was Spectral Biclustering with 3 biclusters (Kluger et al., 2003). Moreover, the ranks of the unconstrained and constrained module arrays were 200 and 74 respectively, which indicates that both arrays were full rank and did not contain redundant modules. This analysis did not result in meaningful biclusters and suggested that there is little relationship between the two types of modules. Nevertheless, the unconstrained modules provided predictions of higher quality than the constrained ones.

**Fig. 13:**
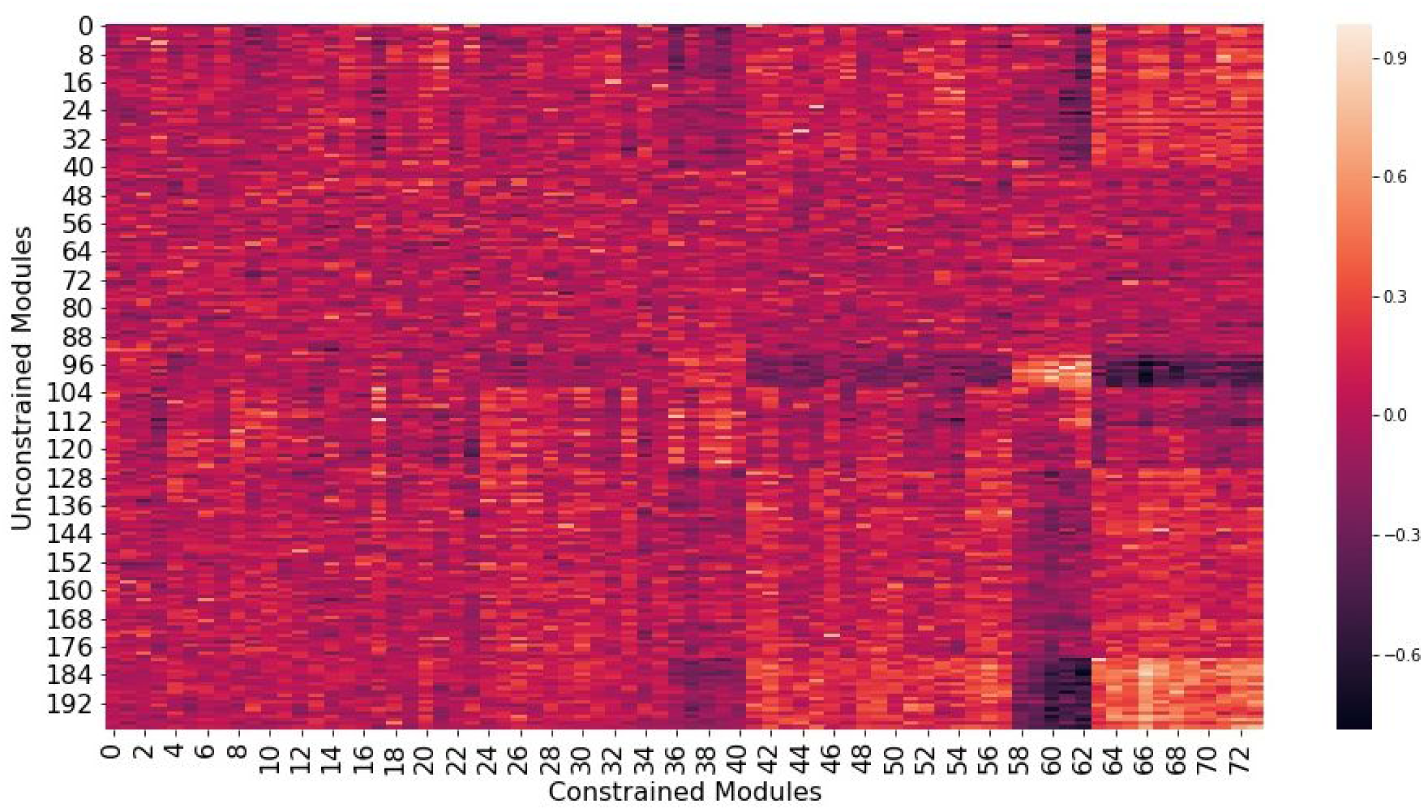
Heatmap displaying correlations between the constrained and unconstrained spatial gene modules. The correlations were estimated with the Spearman’s rank correlation coefficient. 58% out of 6600 correlations in total were considered significant (p ≤ 0.05). However, there are no evident correlation patterns between the two types of modules.

Moreover, gene ontology enrichment analysis was applied to the unconstrained spatial modules and the tract-tracing experiments in order to identify significant annotations related to synaptic and neuronal function in the mouse brain (Rivals et al., 2007). The unconstrained modules were preferred over the constrained ones due to providing better quality predictions. For each tracing experiment, we included the most predictive genes whose coefficients exceeded the 99^*th*^ percentile for that experiment. In the case of modules we included all genes having a non-zero coefficient. The percentage of modules and tracing experiments having at least one significant annotation was 100% and 98% respectively. A tracing experiment was associated with 12 annotations on average (median), while a module was associated with 39 annotations on average.

We observed that annotations related to postsynaptic function were associated with both the RSPagl experiment and module 9 (see figure 14). The jaccard similarity coefficient between the significant annotations of module 9 and modules 70 and 88 was 0.7 and 0.69 respectively, and thus annotations of module 9 were considered to be representative of the three modules.

**Fig. 14:**
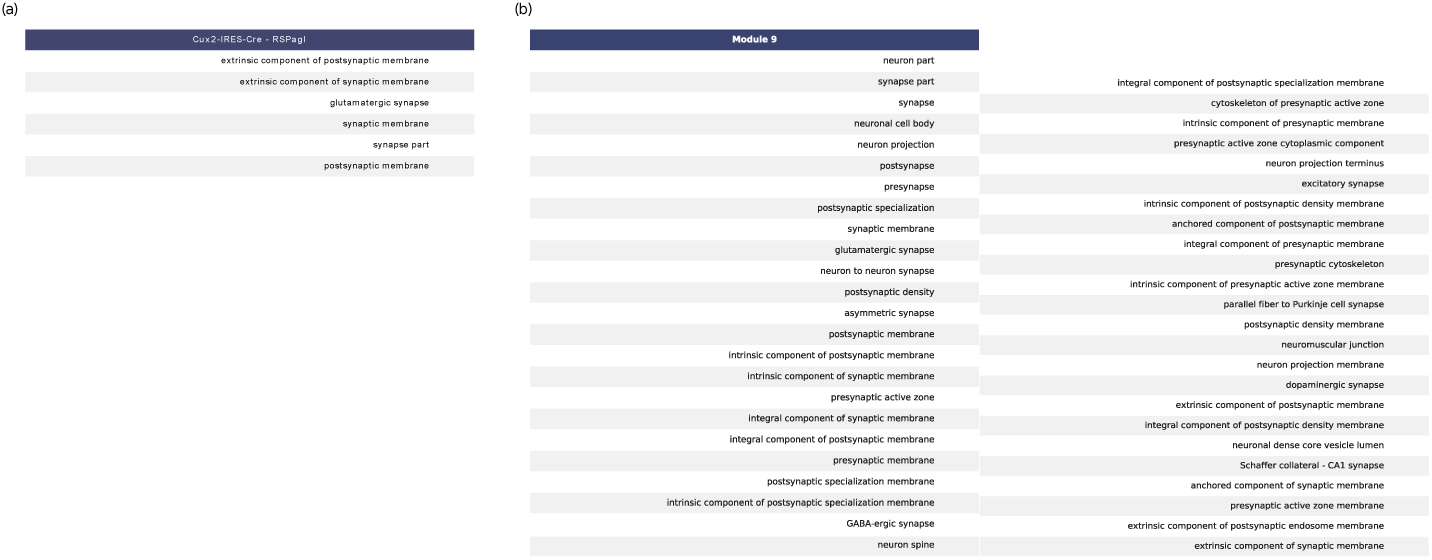
An enrichment analysis reveals annotations of neurons and synapses for a spatial module and a tracing experiment. (a) Significant annotations for a Cux2-IRES-Cre experiment injected in the RSPagl area.

As a generalization of this observation, the percentage of modules and tracing experiments having at least one annotation related to postsynaptic function was 100% and 70% respectively. Hence, annotations with postsynaptic function was another common denominator between a substantial number of tracing experiments and spatial modules, in addition to strong correlations and predictive capability.

## 4 Discussion

In this study we built a predictive workflow, based on ridge regression and random forest based models, to predict axonal projection patterns in the mouse brain using gene expression data. Using the nested k-fold cross-validation technique, we obtained a median *r*^2^ of 0.54 over 1397 tract-tracing experiments. In order to compare with previous studies (Ji et al., 2014), we developed an approach to make binary predictions and obtained similar performance to previous studies. Furthermore, we analyzed the spatial organization of genes in modules defined in different ways, based on the DLSC method, for determining links between gene expression and axonal projection patterns. We obtained median *r*^2^ scores of 0.51 and 0.45 for the unconstrained and constrained approaches respectively. Finally, we applied gene ontology enrichment analysis to gene groups with high coefficient scores, and a substantial number of the groups were found to be associated with annotations related to postsynaptic function. In the following we will put the performance of the different cases in the context of previous studies, interpret our findings, suggest potential future work and discuss strengths, limitations and other applications of our pipeline.

The results of our study are consistent with the findings from the (Ji et al., 2014) study, specifically since our binary approach yielded a similar performance with a median 93% auROC value on wild-type data. In contrast to this study however, we did not rely on arbitrary thresholds for binarizing each tracing experiment to attain a 50% connectivity. Instead, we provided a data-driven estimation of the most optimal threshold value. In addition, we extended their analysis by including cre-line data that had not been subjected to such an analysis before.

When including both cre-line and wild-type data, we found a median auROC value of 0.89 across all 1397 tracing experiments. The increased performance of the models on binary predictions compared to continuous predictions is presumably due to the reduction of projection pattern related information which can therefore be more easily captured by the gene expression data (figure 8). However, binary connectivity descriptions do not inform the modeler about the strength of a projection. Hence, the continuous predictions are more suitable for analytic purposes. For that reason, we provided richer predictions of the mouse mesoconnectome by incorporating continuous patterns to our analysis (figure 6 for continuous predictions and figure 8 for binary ones).

Overall, our ridge regression models provided significant predictions, since they out-performed in every case the surrogate and the null models. This implies that gene expression contains information related to axonal projection patterns in the mouse brain. Regarding the variability of predictions, our statistical tests indicated that the difference in performance between cre-line and wild-type tracing experiments, quantified as *r*^2^ score, was not statistically significant (p = 0.004 for 14000 random permutations). A possible explanation is that both wild-type and cre-line projection patterns fall within the range of predictions that can be covered by the gene expression data. Irrespective of explanation, the results show that the gene expression data contain enough information to also account for the more specific cre-line projection patterns. The ridge regression models trained with spatial gene co-expression modules rather than expressions of individual genes, also outperformed corresponding surrogate and null models (figure 10). However, we found that such predictions were slightly less accurate on average than the gene expression based ones. Despite that, significant predictions of such models and strong correlations between axonal projections and spatial modules suggest that information related to axonal projections is present in modules of genes instead of being present in individual genes.

When comparing constrained modules with the unconstrained ones, we observed dissimilar patterns and an inferior performance for the constrained one when predicting tracing experiments. Such results suggest a lack of direct relation between spatial modules created exclusively by ISH data and modules that were constrained by single cell RNA sequencing data. A possible explanation is that distinct predictive modules were mixed when including all genes differentially expressed in the 74 cell-types, which suggests that better performance could be reached when selecting a subset from amongst them.

Regarding gene ontology enrichment analysis, a substantial number of tracing experiments (70%) and all unconstrained spatial modules (100%) were statistically associated with postsynaptic function. This may suggest that a potential causal link between axonal projections and gene expression in the mouse brain could be gene co-expression modules with a postsynaptic function and specific spatial footprints. This suggestion is consistent with the findings of (Roy et al., 2018), according to which presynaptic and postsynaptic locations have a particular protein profile. These profiles are partially reflected in gene expression data by locally expressed genes at axonal release sites (Glock et al., 2017; Cajigas et al., 2012; Holt and Schuman, 2013). Nevertheless, the causal links are far from being clear and will thus require further work.

A strength of this study was the inclusion of layer and cell-class specific patterns by including cre-line data to our analysis. To our knowledge, this is the first study that predicts brain-wide and cell-class specific projection patterns from gene expression data. Another advantage of this study was that it went beyond solely providing a predictive workflow, and it focused on discovering links between the two data modalities by analyzing the spatial organizations of genes with the dictionary learning and sparse coding technique (Li et al., 2017) and with gene ontology enrichment analysis (Rivals et al., 2007).

We acknowledge some limitations. First, 0.7% of ridge regression based models had an r^2^ score close to zero. This could be attributed to parameters being optimized over all tracing experiments belonging to one cre-line or the wild-type category rather than for each experiment (injection) separately. Therefore, it can be expected that performance will be reduced when multiple projection patterns with a noisy subset need to be predicted simultaneously.

Another explanation could be that including the genetic information of target areas without its relation to source areas has limited capacity in predicting projection patterns. According to (Fulcher and Fornito, 2014), coupled gene expression patterns were shown to be directly linked with the large-scale topology of the mouse mesoconnectome. Furthermore, in (Bleakley et al., 2007) they used the support vector machine algorithm with kernels that coupled the feature vectors of nodes, for inferring the edges of biological networks. As a recommendation for future work, we can adapt this strategy to couple source and target based gene expression patterns and infer their corresponding axonal projections.

Another limitation is that unionization of data leads to information loss, not-a-number values and projection bias because of diversity in sizes of source brain areas. For that reason we will focus our future analyses on the volumetric gene expression and axonal projections data, as to avoid such issues and provide a finer grained predictive pipeline.

Furthermore, ridge regression and random forest based models provided significant predictions of axonal projections from gene expression data, but they are not capable of explicitly modeling the joint distribution between the two data modalities. Such explicit modeling could be advantageous in the case of training models to predict cellular resolved projections since data that could serve as training labels, such as single-neuron axonal reconstruction data, are still limited (Economo et al., 2019; Winnubst et al., 2019). Future directions might include incorporating generative probabilistic models, since models such as the infinite relational model have been successful in capturing the distributions of various connectomes such as the C.elegans connectome and the mouse retina microcircuit (Jonas and Kording, 2015; Ambrosen et al., 2013; Hinne et al., 2014, 2017; Betzel and Bassett, 2017).

Whole brain cellular resolved connections have yet to be described. The capability of our models to provide information for a more faithful reconstruction of the connectome at this resolution will depend on two factors. The first factor will be the ability to incorporate new advances in neuroanatomy and translational neuroscience, such as single-cell RNA sequencing and light sheet fluorescence microscopy (Tasic, 2018; Corsetti et al., 2019; Rolnick and Dyer, 2019).

The second factor will be the ability to mine at a higher spatial resolution from already tested data modalities such as in-situ hybridization based gene expression data. For this factor we will need to adapt additional computational tools for use in our pipeline. One potential tool is spatial point process analysis, which has successfully been used to extract spatially distributed counts of cells and synapses from modalities such as Nissl-stained brain images (LaGrow et al., 2018; Anton-Sanchez et al., 2014).

Translational neuroscientists could benefit from the use of our predictive workflow. A potential use case could include neuroscientists that study the effect of genes in the cognitive processes of the mouse brain. An example gene could be parvalbumin (PV) which according to (Nakazawa et al., 2012) has been linked to schizophrenia. Our workflow can then be used for studying the effect of altering the level and patterns of PV expression on the mesoconnectome and the resulting brain activity, which can in turn by validated by electrophysiological experiments.

Descriptions of potential use-cases for our predictive workflow together with their associated python code can be found online at the HBP Collaboratory and at Github (see table 1). The use cases are intended to be available via the EBRAINS infrastructure, provided as part of the EU-funded Human Brain Project.

Taken together, we have demonstrated a predictive workflow that can further be used to perform multimodal data integration to improve the accuracy of the predicted mouse mesoconnectome using gene expression data and support other neuroscience use cases.

## Acknowledgment

Parts of this research have received funding from the European Union’s Horizon 2020 Framework Programme for Research and Innovation under Grant Agreement No 785907 (Human Brain Project SGA2) and the FLAG ERA project FIIND (NWO054-15-104).

