## supplementary pdf 1 for "Prediction of a cell-type specific mouse mesoconnectome using gene expression data"

### 1 Supplementary Methods

This subsection goes beyond describing methodologies of our predictive workflow and highlights a number of use cases from which potential users of the workflow can benefit.

#### 1.1 Link to Mouse Connectivity Models - Use Cases

Besides integrating our analysis with the Mouse Connectivity Models (MCM) tool (Knox et al., 2018), we implemented and documented use cases that are related to the tool. Specifically, the users can download data with the MCM tool and then obtain the array of interest based on volumetric or regionalized preference. Alternatively, users can give their own projection patterns as input, for instance based on reconstructed axonal projections or predicted ones generated from our models. Such input will be mapped automatically to the volumetric scale of 100 $\mu m^3$ and then provided to the MCM tool for connectivity array construction in a volumetric or regionalized form.

Nestor Timonidis
[1]Neuroinformatics department, Donders Centre for Neuroscience, Radboud University Nijmegen, Heyendaalseweg 135, 6525 AJ Nijmegen, the Netherlands


Rembrandt Bakker
[1]Neuroinformatics department, Donders Centre for Neuroscience, Radboud University Nijmegen, Heyendaalseweg 135, 6525 AJ Nijmegen, the Netherlands
[2]Inst. of Neuroscience and Medicine (INM-6) and Inst. for Advanced Simulation (IAS-6) and JARA BRAIN Inst. I, Jülich Research Centre, Wilhelm-Johnen-Strasse, 52425 Jülich, Germany.

Paul Tiesinga
[1]Neuroinformatics department, Donders Centre for Neuroscience, Radboud University Nijmegen, Heyendaalseweg 135, 6525 AJ Nijmegen, the Netherlands

#### 1.2 Incorporating new user data

The capability of the developed workflow to incorporate external or user-generated data was validated in the form of use cases. A user can get predictions for a new gene expression dataset by selecting a layer and class profile (i.e. L5 CT corresponding to layer 5 corticothalamic specific projections) and a source area of interest (i.e. MOp or primary motor area). The model most closely associated with the selected preferences will automatically be selected and the predicted axonal projections will be returned. Moreover, a user can decide whether to trust the selected model based on its performance score ($r^2$) or select another model. Furthermore, the patterns can be converted to a laminar specific regionalized connectivity matrix based on the MCM tool as described in subsection 1.1. Finally, the visualization part of our workflow can be used for a visual inspection of the various projection patterns (subsection 1.3).

#### 1.3 Brain Visualization

Part of our predictive worfklow comprised visualizations of brain volumetric data in the form of cortical flatmaps and brain slices. We constructed both cortical flatmaps oriented along the anterior-posterior and left-right axes, and brain slices oriented along the inferior-superior and left-right axes. The data were then converted to either JavaScript Object Notation (JSON) format or in Neuroimaging Informatics Technology Initiative (NIfTI) format, in order to be visualized through an API call to the Scalable Brain Atlas (SBA) Composer, a 3D brain visualization tool (Bakker et al., 2015). The SBA composer provides 3-dimensional visualizations of brain volumes through a user-friendly interface.
In order to appreciate the spatial context of our predicted data, they were mapped to a 25 or 10 $\mu m^3$ volumetric scale based on annotation volumes. Afterwards, they were plotted overlaid with the templates that were provided together with the annotation volumes by the Allen Institute (table 1). To achieve this we used the red, green and blue channels to represent the components as images and then mixed them by selecting per pixel the component with the highest intensity. More details about the aforementioned formats and tools can be found in table 1.

### 2 Supplementary Figures

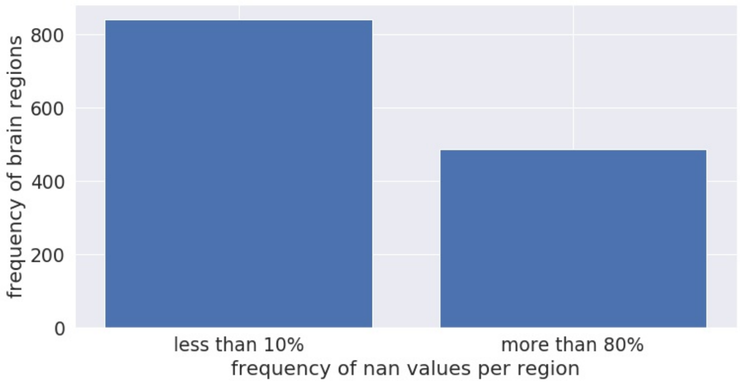


Fig. 1: Histogram with frequency of NaN value occurrence per brain area. Gene Expression dataset.

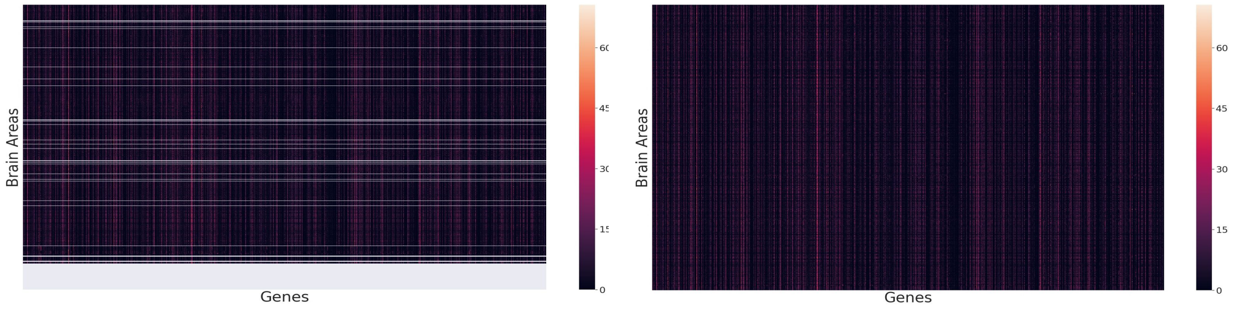


Fig. 2: Gene Expression (left panel) before and after (right panel) NaN removal

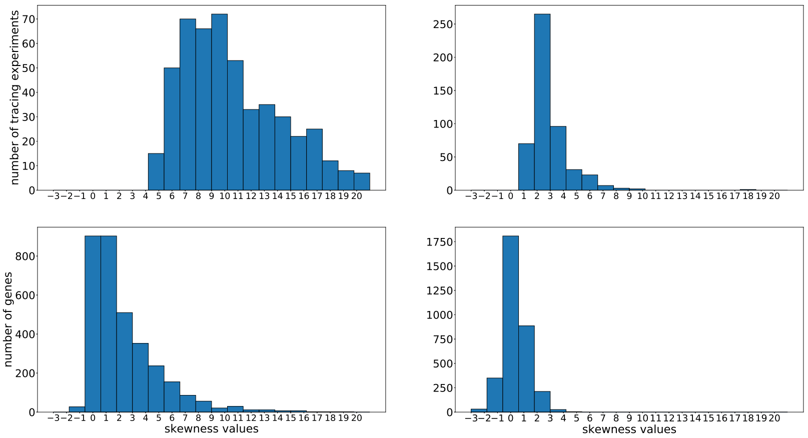


Fig. 3: Skewness distributions for wild-type projection strength (top) and gene expression (bottom) datasets before pre-processing (left panels) and after pre-processing (right panels).

Table 1: Hyperlinks for websites, tool descriptions and format descriptions related to our analysis. See main text for details.

| | |
|---|---|
| Allen Institute | `https://alleninstitute.org/` |
| CC documentation | `https://allensdk.readthedocs.io/en/latest/connectivity.html` |
| CCF v3.0 | `http://help.brain-map.org/display/mouseconnectivity/Documentation` |
| MCC use case | `https://alleninstitute.github.io/AllenSDK/_static/examples/nb/mouse_connectivity.html` |
| NIfTI | `https://nifti.nimh.nih.gov/` |
| JSON | `https://en.wikipedia.org/wiki/JSON` |
| SBA | `https://scalablebrainatlas.incf.org/composer-dev/?template=ABA_v3` |
| Bioconductor software | `http://bioconductor.org/packages/release/data/annotation/html/org.Mm.eg.db.html` |
| Repository of our Code on the HBP Collaboratory | `https://collab.humanbrainproject.eu/#/collab/8650/nav/65518` |
| Repository of our Code on Github | `https://github.com/ntimonid/Connectomic-Composition-Predictor-CCP-` |
